# Single-cell multi-scale footprinting reveals the modular organization of DNA regulatory elements

**DOI:** 10.1101/2023.03.28.533945

**Authors:** Yan Hu, Sai Ma, Vinay K. Kartha, Fabiana M. Duarte, Max Horlbeck, Ruochi Zhang, Rojesh Shrestha, Ajay Labade, Heidi Kletzien, Alia Meliki, Andrew Castillo, Neva Durand, Eugenio Mattei, Lauren J. Anderson, Tristan Tay, Andrew S. Earl, Noam Shoresh, Charles B. Epstein, Amy Wagers, Jason D. Buenrostro

## Abstract

*Cis*-regulatory elements control gene expression and are dynamic in their structure, reflecting changes to the composition of diverse effector proteins over time^1–3^. Here we sought to connect the structural changes at *cis*-regulatory elements to alterations in cellular fate and function. To do this we developed PRINT, a computational method that uses deep learning to correct sequence bias in chromatin accessibility data and identifies multi-scale footprints of DNA-protein interactions. We find that multi-scale footprints enable more accurate inference of TF and nucleosome binding. Using PRINT with single-cell multi-omics, we discover wide-spread changes to the structure and function of candidate *cis*-regulatory elements (cCREs) across hematopoiesis, wherein nucleosomes slide, expose DNA for TF binding, and promote gene expression. Activity segmentation using the co-variance across cell states identifies “sub-cCREs” as modular cCRE subunits of regulatory DNA. We apply this single-cell and PRINT approach to characterize the age-associated alterations to cCREs within hematopoietic stem cells (HSCs). Remarkably, we find a spectrum of aging alterations among HSCs corresponding to a global gain of sub-cCRE activity while preserving cCRE accessibility. Collectively, we reveal the functional importance of cCRE structure across cell states, highlighting changes to gene regulation at single-cell and single-base-pair resolution.

## Introduction

Through homeostasis, development, and disease, *cis*-regulatory elements change in structure and recruit new regulatory proteins, which define the overall function of the element^4^. In this process, *cis*-regulatory elements act as hubs of gene regulation to establish primed, activated, or repressed genes and determine the overall function and potency of cells^1, 3, 5, 6^. These structural changes are largely mediated by the competition of nucleosomes, TFs and transcriptional machinery, which dynamically slide, evict and recruit effector proteins^7, 8^. Despite this rich understanding of the biochemical activities occurring on DNA, in genomics individual *cis-* regulatory elements are often studied as discrete functional units, motivating a need for genomic tools that trace chromatin structure at single-base-pair resolution.

Methods that measure chromatin accessibility have revealed a diverse repertoire of cCREs^6^. Additionally, DNA footprinting methods elucidate TF binding at cCREs by quantifying the protection of DNA from chemical^9^ or enzymatic^10–13^ cleavage, yielding base-pair resolved maps of diverse proteins bound to DNA^14^. Using high-throughput DNA sequencing, footprinting is now performed genome-wide, revealing the function of non-coding genetic variation^15^ and improving the construction of gene regulatory networks^16–18^. However, despite best efforts, footprinting methods are afflicted with sequence bias severely limiting accuracy^19^. Further, computational methods for footprinting are optimized to detect binding of TFs of typical size, excluding the analysis of nucleosomes or atypical TFs. These limitations preclude our ability to measure intra-cCRE structural dynamics that reflect changes to the composition and function of cCREs over time.

Here, we develop PRINT (<u>P</u>rotein-<u>R</u>egulatory element Interactions at <u>N</u>ucleotide resolution using <u>T</u>ransposition), a framework that i) accurately corrects for sequence bias in chromatin accessibility data, ii) computes the interaction of DNA with objects of various sizes (multi-scale footprinting) and iii) leverages single-cell multi-omics to identify the structural changes to cCREs and their impact on gene expression. Using this approach, we show that DNA bound proteins, including TFs and nucleosomes, create unique cleavage patterns and demonstrate that multi-scale footprints enable accurate prediction of TF binding genome-wide. Next, we combine multi-scale footprinting with single-cell multi-omic data across human hematopoiesis to track TF and nucleosome binding dynamics across differentiation. We discover wide-spread restructuring of cCREs during differentiation, wherein nucleosomes slide, expose new sites for TF binding and promote gene expression. We refer to the genomic regions that modularly expand and shrink within cCREs as “sub-cCREs” and show that sub-cCREs can explain changes to gene expression in the absence of overt changes to chromatin accessibility. Finally, we define sub-cCRE dynamics in response to aging of mouse hematopoietic stem cells (HSCs). Here, we find that many age-associated promoter alterations increase TF binding of sub-cCREs, while maintaining the overall accessibility of the element. Overall, we find sub-CREs as regulators of gene expression and cell state, revealing a unique structure of gene regulation at single-cell and single-base-pair resolution.

### Multi-scale footprinting enables detection of DNA binding by factors of diverse sizes

To enable tracking of structural changes within cCREs we developed PRINT, a computational approach to footprint proteins of diverse sizes (**Fig. 1a**). To do this, we sought to use bulk or single-cell ATAC-seq data as input. However, Tn5 transposase has a strong sequence preference^13, 20^, which may significantly confound footprint detection. To evaluate and create approaches for modeling Tn5 sequence bias, we generated high-coverage Tn5 insertion data on deproteinized DNA from bacterial artificial chromosomes (BACs) containing a total of 5.6 Mb of the human genome (**Extended Data Table 1**). A total of 193.2 million reads aligned to the BACs, resulting in 34.5 Tn5 insertions per base-pair. We also performed 5 biological replicates and found that the observed Tn5 bias is highly reproducible (R > 0.97, **Extended Data Fig. S1a-c**).

**Figure 1.**
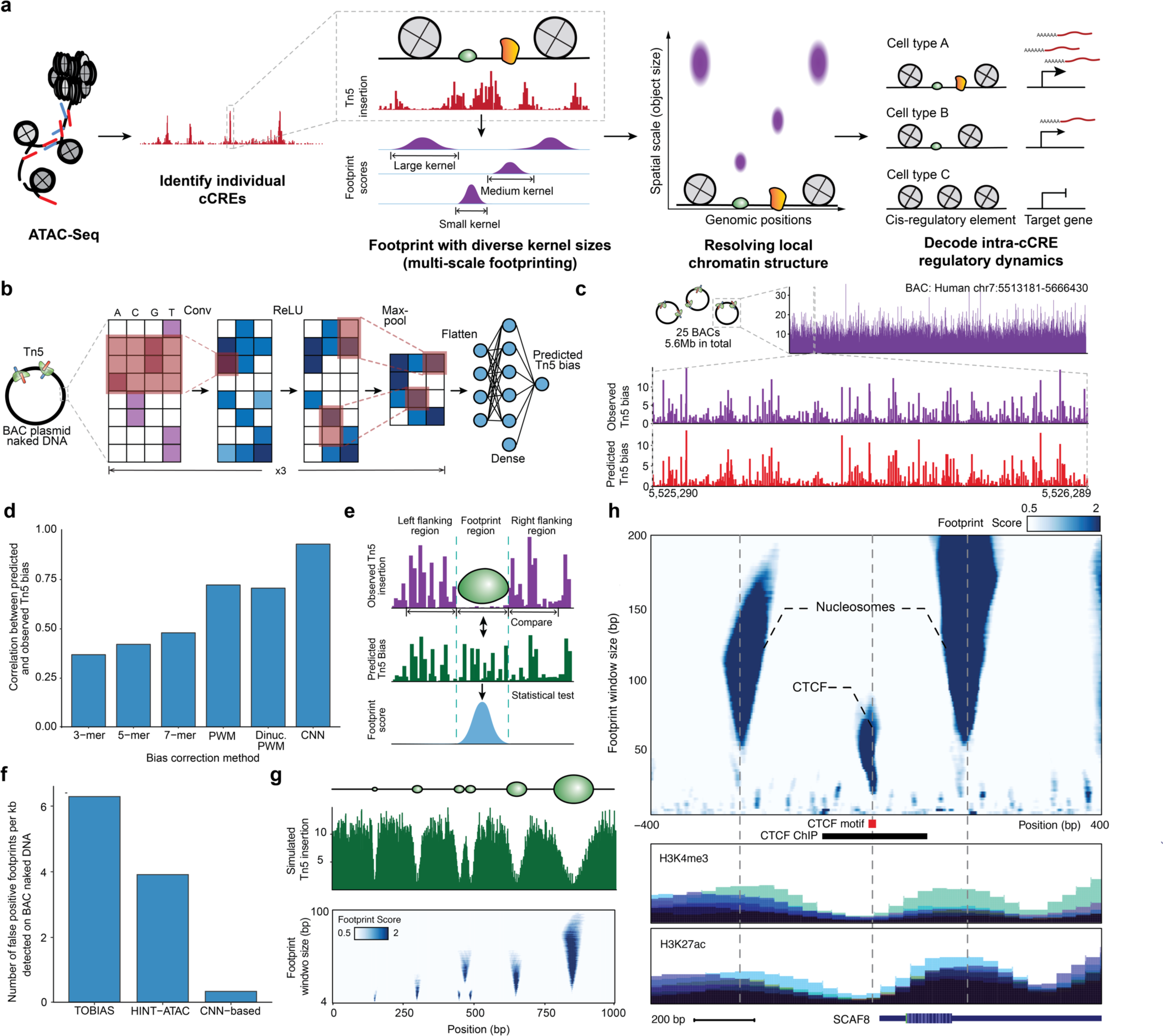
Multi-scale footprinting detects DNA-protein interactions at various spatial scales. **a**, Overview of the multi-scale footprinting workflow. **b**, Schematic illustration of the Tn5 bias prediction model. **c**, Single-nucleotide resolution tracks of observed and predicted Tn5 bias on naked DNA in the BAC RP11-93G19. **d**, Bar plot comparing performance of the CNN model with previous bias correction models. **e**, Schematic illustration of footprint score calculation. **f**, Bar plot showing the frequency of calling false positive footprints by previous ATAC-footprinting methods and our method. **g**, Multi-scale footprints with simulated objects. Top: schematic of simulated objects with various sizes. Middle: Simulated single base pair resolution Tn5 insertion tracks based on the above objects. Bottom: Heatmap showing the multi-scale footprints calculated based on the simulated Tn5 insertions. The horizontal axis represents single base pair positions, and the vertical axis represents footprint window sizes. **h**, Multi-scale footprints in the cCRE region chr6:154732871-154733870. Bottom tracks are histone ChIP signals obtained from ENCODE.

Using the BAC data, we trained a convolutional neural network that takes as input DNA sequence and predicts Tn5 sequence preference (**Fig. 1b**). We found that deep learning achieved a correlation of 0.94 between predicted and observed bias, significantly outperforming k-mer and PWM models (**Fig. 1c, d**) while achieving the highest improvements in regions of high GC-content (**Extended Data Fig. S1d,e**). Exemplifying the utility of modeling Tn5 preference, we provide Tn5 bias prediction for the entire human genome, alongside common model organisms including *Pan troglodytes, Mus musculus*, *Drosophila melanogaster*, *Saccharomyces cerevisiae*, *Caenorhabditis elegans* and *Danio rerio,* covering a total of ∼11B bases of DNA sequence. We also provide a pre-trained deep learning model that can be extended to any new species or applied to personal genomes (see **Data Availability**).

To call footprints, we developed a statistical approach for footprinting that quantifies the depletion of observed Tn5 insertions relative to the Tn5 sequence bias, resulting in a footprint score representing the statistical significance (-log10 *p*-value) for each base pair position (**Fig. 1e**, Methods). Using our deproteinized BAC data as a control, we detected little to no footprint signal using our approach on naked DNA (**Extended Data Fig. S1f-o)**. In contrast, prior footprinting methods^21, 22^ report up to 35,262 false positive footprints within the 5.6 Mb BAC regions, corresponding to an average false positive rate of 23% across all TFs (**Fig. 1f**). The Tn5 bias model and statistical approach described here reduced the number of false positive footprints by approximately one order of magnitude (**Fig. 1f**), demonstrating that bias correction is essential for accurate footprinting.

Finally, we explored footprinting across spatial scales to detect DNA-bound proteins of different sizes. We performed footprinting, using simulated data (**Fig. 1g**) and ATAC-seq data (**Fig. 1h**), with window sizes ranging between 4-200 base pairs and observed drastically different footprint patterns corresponding to TF and nucleosomes (**Fig. 1h**). Therefore, we reasoned that multi-scale footprinting may fractionate molecular interactions at different scales and outline the local physical structure of chromatin.

### TFs and nucleosomes have signature multi-scale footprints

Inspired by the diversity of structures seen in multi-scale footprints, we sought to categorize proteins by footprint sizes and shapes. We obtained TF ChIP-seq data^6^ and generated aggregate multi-scale footprints to find that TFs may leave small (CREB1) and large (CTCF) footprints (**Fig. 2a-b**, **Extended Data Fig. S2a-c**). We found that TFs (n = 112) clustered into 6 distinct groups based on their size, shape, and footprint strength (**Extended Data Fig. S2d**). We found that the majority of TFs in cluster 1, 4, and 5 leave visible footprints (n = 71) at 20 bp and 40 bp scales, whereas TFs in cluster 2, 3, and 6 (n = 41) leave weak or no footprints at these same size scales. We also validated that footprints of larger sizes (100-140bp) correspond to prior measures of nucleosome position^23, 24^ (**Fig. 2c**). In summary, multi-scale footprints reveal diverse DNA-protein interactions, enabling the analysis of both TFs and nucleosomes in one computational approach.

**Figure 2.**
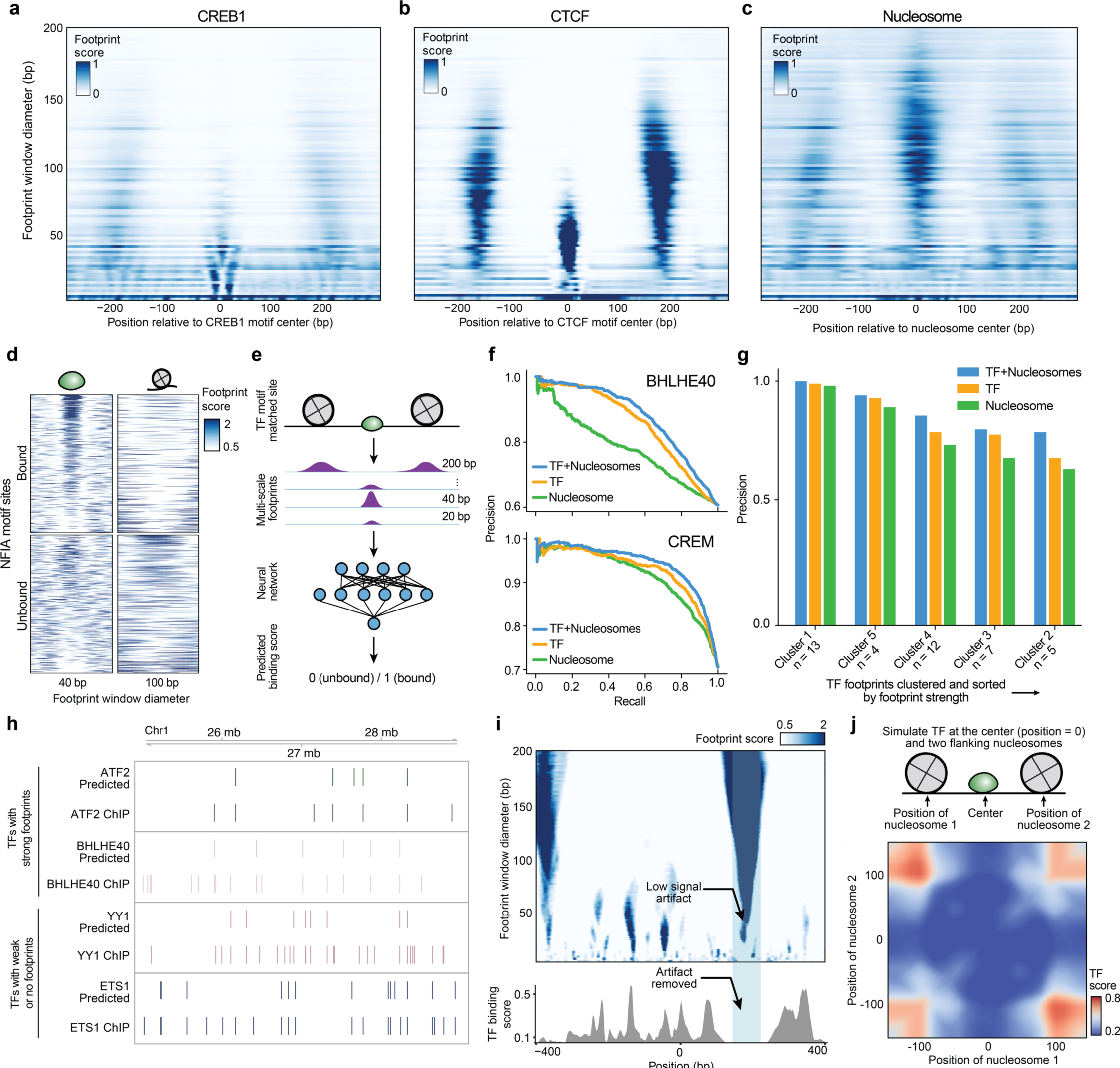
TFs and nucleosomes binding leave signature multi-scale footprint patterns. **a,b**, Multi-scale aggregate footprints for TFs CREB1 and CTCF. The x-axis represents the position relative to the center of the TF motif, and the y-axis represents footprint scores computed using each footprint window size. **c**, Multi-scale aggregate footprints for nucleosomes. The x-axis represents the position relative to the center of the nucleosome as determined by chemical mapping, and the y-axis represents footprint scores computed using each footprint window size. **d**, Multi-scale footprints around individual bound and unbound NFIA motif sites. Each row represents a single locus with a matched NFIA motif. **e**, Schematic illustration of training TF binding prediction models using multi-scale footprints as input. **f-g** Ablation test results. **f**, Example precision-recall curves of cluster 1-specific models trained without masking, with TF masking, and with nucleosome masking, respectively. **g**, Bar plot showing precision of the TF habitation model when trained without masking, with TF masking, and with nucleosome masking, respectively. **h**, Comparison between predicted and ChIP-detected TF binding sites. Only sites with a matched TF motif are included. **i**, Top: heatmap showing multi-scale footprints within the cCRE at chr11:67629937-67630936. The x-axis represents single base pair positions in the cCRE, and the y-axis represents footprint window size. Bottom: predicted TF binding scores within the same region. **j**, Heatmap showing predicted TF habitation score for different simulated TF and nucleosome configurations. Horizontal and vertical axes represent the distances of the two simulated nucleosomes from the center TF.

Motivated by these observations, we trained a neural network classifier that uses multi-scale footprints and motif positions as input to predict TF binding (**Fig. 2d,e** and **Extended Data Fig. S3a-c**). Provided that cluster 1 TFs leave the strongest footprints, we first trained the model predicting TF binding of cluster 1 TFs using multiscale footprints. The model achieved a median precision of 0.71 on held out test ChIP data^25^, outperforming prior methods (0.65 for HINT-ATAC and 0.62 for TOBIAS when benchmarked at a matched recall, **Fig. 2f** and **Extended Data Table 2**).

We next sought to extend this approach to TFs that leave weak or undetectable footprints. We trained a new model using data from all 6 clusters of TFs. As many TFs (37%) do not leave clear footprints, this model further prioritizes nucleosome position for TF binding prediction. As such we refer to this model as the “TF habitation model” and its prediction scores as “TF habitation scores”. The TF habitation model achieved a median precision of 0.76 for cluster 1 TFs and 0.67 across all TFs on held out K562 data (n = 41), again outperforming previous methods (0.58 for HINT-ATAC and 0.59 for TOBIAS, **Extended Data Fig. S3d,e**). We next tested model performance on primary cell samples, expanding the analysis to 91 TF binding data datasets in total^6^. The model achieved a median precision of 0.73 across all TFs while recovering hundreds to thousands of binding sites per TF (**Fig. 2g-h** and **Extended Data Table 2**). Additionally, we determined a 0.8% false positive rate using the BAC data. In conclusion, using multi-scale footprinting, we developed an approach that accurately predicts protein-DNA interactions at multiple length scales.

Upon further investigation we find that these models use nearby nucleosomes, together with TF footprints, to improve predictions (**Fig. 2i, j**). As expected, the model uses a high TF footprint signal (40 bp) at the motif center and low nucleosome signal (100-200 bp) surrounding the motif (**Extended Data Fig. S3f**) for prediction. Additionally, we found frequent cases wherein nucleosome footprints span lower (40bp) scales, but are correctly identified as artifacts by the model (**Fig. 2i**). Interestingly, the model also found that high nucleosome signal distal to the motif (∼100-150 bp) is predictive of TF binding. To further explore the model, we simulated TF and nucleosome footprint within a 300 bp window to find that TF binding scores significantly decrease as nucleosomes approach the TF motif (**Fig. 2j**) or as nucleosomes become delocalized or “fuzzy” (**Extended Data Fig. S3g**). To quantitatively assess improvements, we performed ablation tests wherein TF or nucleosome footprints are removed during training (**Fig. 2f, g**). Using this approach, we observed decreased precision after ablating nucleosomes and found that nucleosomes, without TF footprints, may be highly predictive of TF binding (e.g., CREM). Altogether this indicates that nucleosome position strongly influences TF binding.

### Emerging modular structures of intra-cCRE dynamics

We reasoned that a single-cell multi-omic analysis of footprints would enable pseudo-time-resolved tracking of protein-binding and connect these changes to alterations in gene expression. To generate multi-omic data at a throughput and depth needed for footprinting, we used SHARE-seq^26^ (ATAC and RNA) to profile 874,480 total cells from 7 human bone marrow donors. The resulting data represents a total of 935,959,306 nuclear ATAC fragments and 608,148,224 RNA UMIs across all major hematopoietic cell types, including hematopoietic stem cells (HSCs) and differentiated cell types (**Fig. 3a**, **Extended Data Fig. S4a-e**).

**Figure 3.**
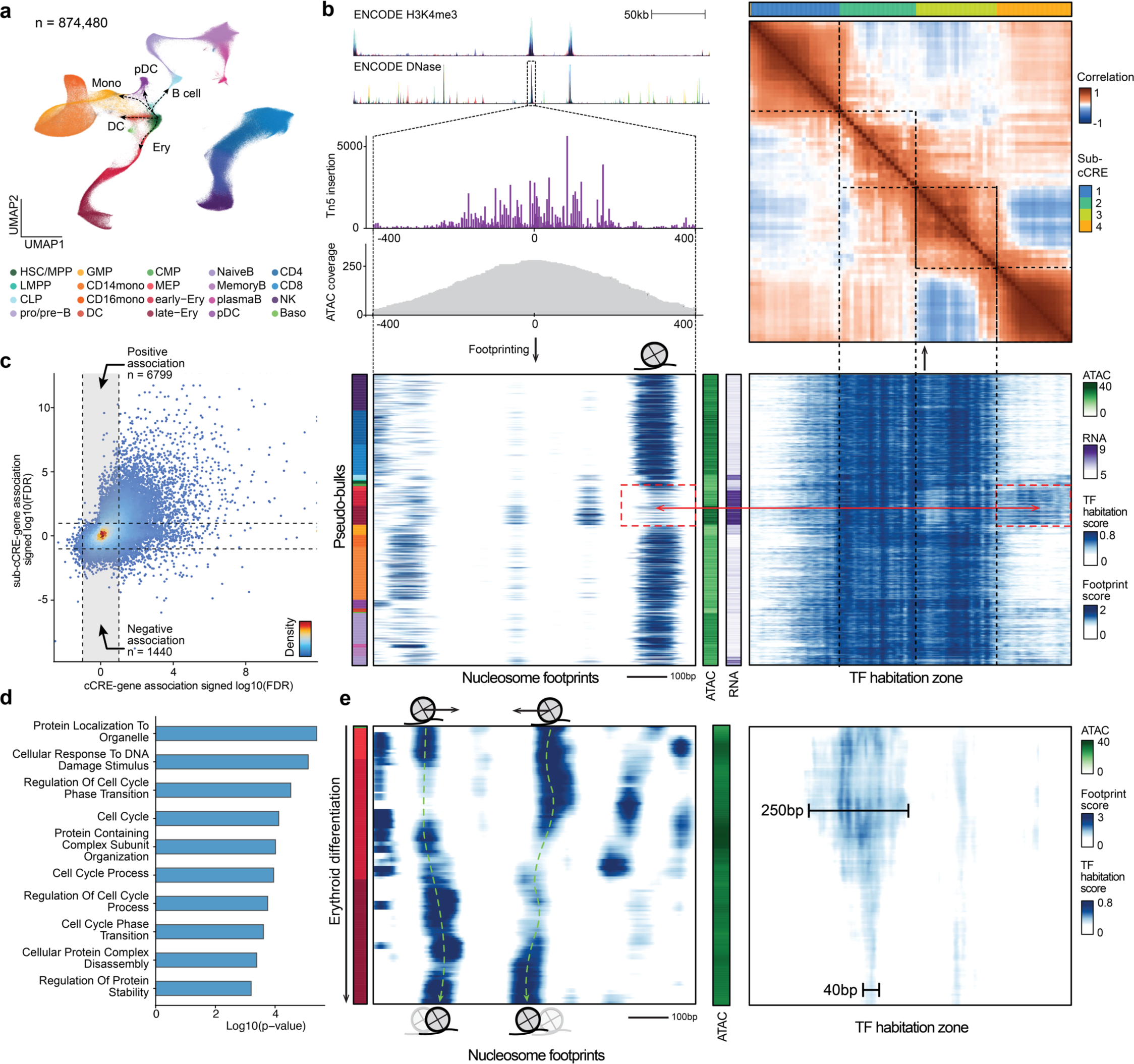
Emerging modular structures of intra-cCRE dynamics. **a**, UMAP of the human bone marrow SHARE-seq dataset. **b**, Example of sub-cCREs. Top left: Tracks showing chromatin accessibility and single-base pair resolution Tn5 insertion in the cCRE at chr4:173334022-173335021. Bottom left and bottom right: Heatmap of nucleosome footprints (100 bp scale) and TF habitation scores in the same region across all pseudo-bulks, respectively. Each row corresponds to a single pseudo-bulk, while each column represents a single base pair position in the cCRE. Left color bar shows the cell type annotation of each pseudo-bulk. Color palette is the same as in **a**. Middle color bars show total accessibility within the cCRE and RNA level of the gene *HMGB2* in each pseudo-bulk, respectively. Top right: Heatmap showing correlation of TF habitation scores between any two positions within the cCRE. Top color bar shows results of automatic segmentation of the cCRE into sub-cCREs. **c**, Scatter plot comparing cCRE-gene correlation and sub-cCRE-gene correlation. For each cCRE, the sub-cCRE with the strongest correlation is selected. Dashed lines represent the FDR threshold of 0.1. **d**, Bar plot showing pathway enrichment of genes with significant sub-cCRE-gene correlation but not cCRE-gene correlation (FDR < 0.1). **e**, Nucleosome tracking across erythroid differentiation. Left: Heatmap of nucleosome footprints in the region chr7:99471434-99472433 across pseudo-bulks in the erythroid lineage. Pseudo-bulks are ordered by pseudo-time. Right: heatmap of TF habitation scores in the same region and pseudo-bulks. Left color bar shows the cell type annotation of each pseudo-bulk. Color palette is the same as in **a.** Middle color bar shows total accessibility within the cCRE in each pseudo-bulk.

Using these single-cell data we sought to define the dynamics at a sub-cCRE scale using PRINT. We generated 1,000 pseudo-bulks encompassing all major cell types and major developmental transitions (**Extended Data Fig. S4f-l**, Methods). Next, we applied multi-scale footprinting and our TF habitation model to these hematopoietic pseudo-bulks. Within individual cCREs, we observed modular structures reflecting gain or loss of TF habitation scores across pseudo-bulks, which we refer to as “sub-cCREs” (**Fig 3b**, **Extended Data Fig. S5a,b**). To quantitatively detect such sub-cCREs, we computed the correlation of TF habitation scores between all positions within each cCRE. The results again show modular structures within the cCRE, as exemplified by regions showing strong off-diagonal correlation (**Fig. 3b**, top-right panel, **Extended Data Fig. S5c,d**). Using such intra-cCRE correlation maps as input, we designed an algorithm to segment each cCRE into sub-cCREs with strong self-association. As a result, we detected 265,070 sub-cCREs across human hematopoiesis. We observed a positive association between cCRE accessibility and the number of sub-cCREs detected inside the cCRE. For the top 10,000 accessible cCREs, we detected on average 3.7 sub-cCREs within each cCRE and the average size of a sub-cCREs is 211.9 bp, which is approximately the size of a nucleosome flanked with linker DNA. As a result, cCREs do not appear to have a fixed boundary or size, but instead shrink, expand, merge or split as modular sub-cCREs lose or gain activity across differentiation (**Extended Data Fig. S5e,f**).

We next sought to examine if the activity of independent sub-cCREs is associated with gene expression variation across cell types. For every cCRE, we computed the correlations between the accessibility of the cCRE and the RNA levels of nearby genes (+/-50 kb). We next computed correlations between the activity of each sub-cCRE (as defined by the average TF habitation score with the sub-cCRE) within this cCRE with the same nearby genes. Interestingly, cCRE-gene and sub-cCRE-gene correlations show divergent association (**Fig. 3c**). A total of 8,239 sub-cCREs were significantly correlated to gene-expression while the corresponding cCREs were not (permutation test, FDR < 0.1). In these cases we find that cCREs re-organize TF binding while maintaining overall accessibility of the cCRE (**Fig. 3b**, bottom panels, **Extended Data Fig. S5g,h**). Furthermore, at these regions we observed strong enrichment in pathways related to cell cycle, proteostasis, and DNA damage response, suggesting a unique mode of regulation for such pathways (**Fig. 3d, Extended Data Table 3**).

Sub-cCREs are approximately ∼200 bp in size, similar to the size of nucleosomes plus linker regions. Hence, we hypothesized that the activity of sub-cCREs is driven by the dynamics of nucleosomes. To further explore this idea, we tracked nucleosome positioning and sub-cCRE dynamics across pseudo-time along erythroid differentiation. We observed nucleosome dynamics in the form of binding, eviction, as well as sliding accompanied by sub-cCRE activation/repression at the same locus (**Fig. 3e**, **Extended Data Fig. S5i,j**), providing evidence for nucleosome reorganization during native human hematopoiesis.

### Intra-cCRE dynamics in hematopoietic aging

Aging is a major risk factor for many highly prevalent diseases such as cancer, cardiovascular disease and neurodegeneration^27^. Extensive previous studies have shown that aging is accompanied by widespread “epigenetic decline”^28–30^. In particular, HSCs have been shown to be compromised in function during aging, contributing to deficient pathogen-and vaccine-evoked immunity and heightened inflammatory responses^31–33^. Mutation of genes involved in epigenetic and chromatin remodeling has frequently been observed in humans with clonal hematopoiesis of indeterminate potential (CHIP), an age-associated condition characterized by the expansion of somatically mutated hematopoietic cell clones, a process associated with an increased risk of hematopoietic malignancy, cardiovascular disease, stroke and all-cause mortality^34, 35^. As such, prior studies have investigated alterations to DNA methylation, heterochromatin or chromatin accessibility during HSC aging^36^. Here, we hypothesized that aging HSCs relocalize regulatory proteins to restructure sub-cCREs and alter the expression of aging genes. Thus, we applied PRINT to discover alterations of TF and nucleosome binding in young or aged hematopoietic cells.

We isolated hematopoietic progenitor cells (Lineage^-^) and HSCs (Lineage^-^ Sca-1^+^ c-Kit^+^ Cd48^-^ CD150^+^) from the bone marrow of young (11 weeks old, n = 10) or aged (24 months old, n = 5) mice by FACS. We then obtained joint ATAC-RNA profiling using the 10x platform (**Fig. 4a**, **Extended Data Fig. S6a-c**, Methods). Consistent with previous studies, we observed an expansion of the HSC compartment during aging^33^ (**Extended Data Fig. S6d**). After QC filtering, we obtained 48,225 cells covering 14,640 HSCs and 33,585 hematopoietic progenitor cells in the mouse bone marrow (**Fig. 4b**, **Extended Data Fig. S7a**). From the Lineage^-^ single-cell data we confirmed an age-associated increase in HSC frequency (**Extended Data Fig. S7b**). Further, HSCs clustered into 3 clusters reflecting points along a continuum of age-related cell states (**Fig. 4c, d**). Validating these clusters, we used gene expression to find age-specific marker genes (*Nupr1*, *Clu*, *Selp*) (**Extended Data Fig. S7c-e**), consistent with findings of previous work^36, 37^. Of the two aging clusters, we found that cluster 6 was more similar to young HSCs (R = 0.95 vs R = 0.91). We therefore refer to the three clusters as young, young-like old, and old HSC states (**Fig. 4d**). Focused on age-related alterations, we used the ATAC profile of single cells to define pseudo-bulks^38^ revealing representative cell states (**Fig. 4e, f**, **Extended Data Fig. S7f**).

**Figure 4.**
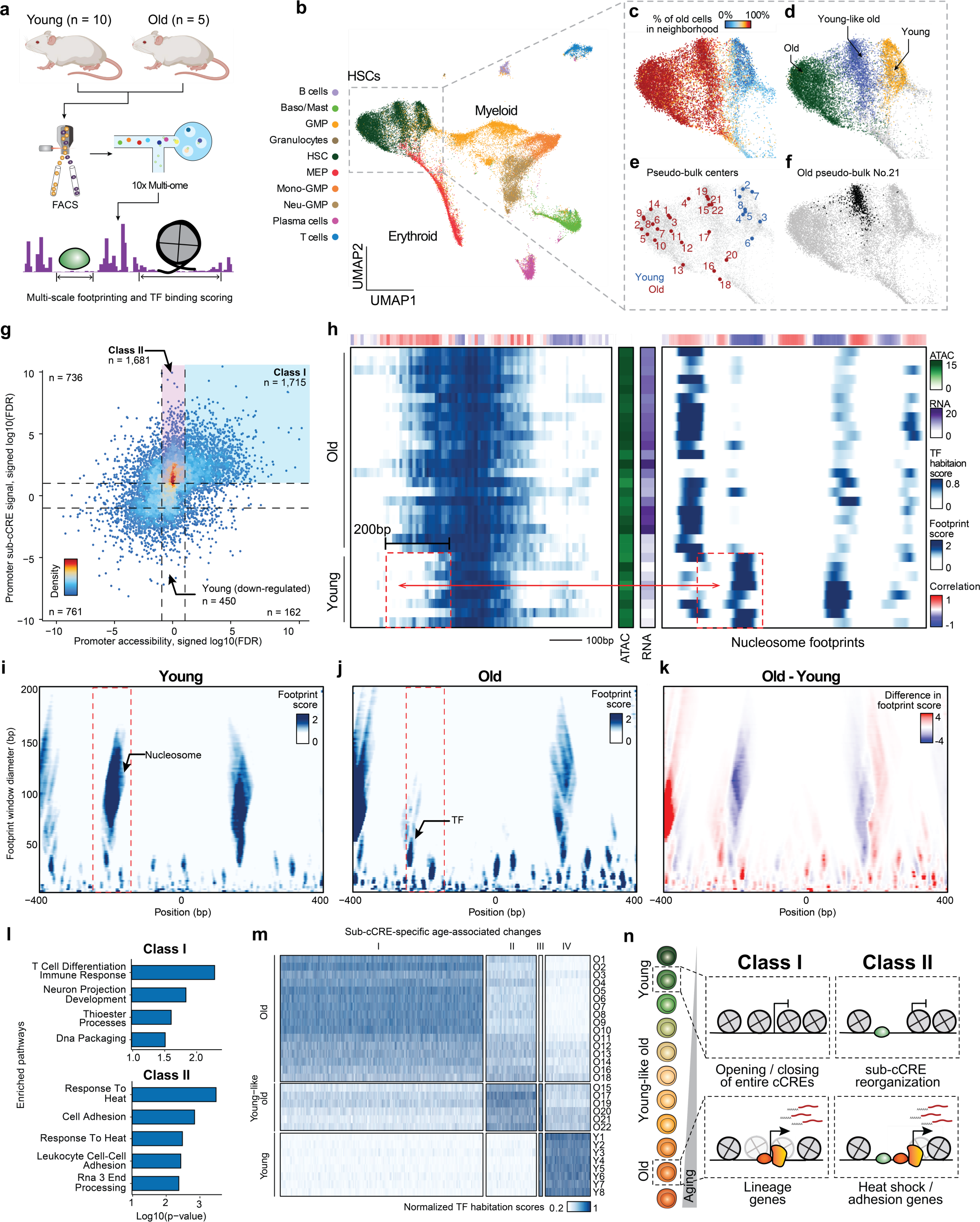
Intra-cCRE dynamics in hematopoietic aging. **a**, Schematic illustration of dataset generation and analysis. **b-f,** UMAP of HSC and progenitor cells. **b**, Cell type annotation. **c**, Percentage of old cells in the 100-cell nearest neighborhood. **d**, Young, young-like old, and old HSC clusters. **e**, Representative cell states detected by SEACells. **f**, Example pseudo-bulk. Black dots represent member cells in old pseudo-bulk 21. **g**, Scatter plot comparing differential cCRE testing and differential sub-cCRE testing results for promoters of differentially expressed genes. Dashed lines represent the FDR = 0.1 threshold. **h**, Heatmaps of TF habitation scores and nucleosome footprints (100 bp scale) within the promoter of *Cdc25b* at chr2:131186436-131187435. Each row corresponds to a single pseudo-bulk, while each column represents a single base pair position in the cCRE. Middle color bars show total accessibility within the cCRE and RNA of *Cdc25b* in each pseudo-bulk, respectively. **i-k**, Heatmaps showing the multi-scale footprints within the *Cdc25b* promoter across age groups. The horizontal axis represents single base pair positions, and the vertical axis represents footprint window sizes. **i**, Young. **j**, Old. **k**, difference between young and old. **l**, Bar plot of pathway enrichment Amy using either Class I or Class II as foreground and the other category as background. **m**, Heatmap showing activity of age-related differential sub-cCREs across pseudo-bulks. Rows correspond to pseudo-bulks and columns represent sub-cCREs. **n**, Schematic illustrating contrasting two classes of age-related cCRE changes (modulation of overall cCRE accessibility and intra-cCRE reorganization).

We first applied PRINT to examine intra-cCRE reorganization in promoters of genes expressed in an age-variant manner. More specifically, we identified promoters of genes with differential expression (**Extended Data Fig. S7e**, **Extended Data Table 4**), and segmented these promoters into sub-cCREs. Applying differential testing among young and old pseudo-bulks, we detected 4,132 old-specific and 1,373 young-specific sub-cCREs (**Fig. 4g**, two-sample t test, FDR < 0.1). We identified age-associated elements with robust increase to the overall accessibility and sub-cCRE activity (“Class I”; n = 1,715). In contrast, we discovered age-associated increases to sub-cCRE activity that are missed when assessing the overall accessibility of cCREs (“Class II”; n=1,681). As one example of Class II promoters, in young cells the promoter of *Cdc25b* is flanked by two nucleosomes. In aged cells, the −1 phased nucleosome is lost, exposing additional DNA for TF binding, which is accompanied by increased gene expression (**Fig. 4h-k**). Strikingly, promoters showing age-related changes only at the sub-cCRE level (i.e., Class II promoters) were enriched for genes in the heat shock pathway, such as *Hsp90ab1*, *Dnaja2* (*Hsp40* member), *Bcl2l11*, and *Ubqln2*, suggesting dysregulated proteostasis during aging (**Fig. 4l**, **Extended Data Fig. S8a**, **Extended Data Table 5**). This suggests that intra-cCRE reorganization might be involved in age-related impairment of proteostasis as reported by previous studies in model organisms^39–41^ and in HSCs^42^. Additionally, we observed enrichment in pathways related to cell adhesion, involving genes such as *Igf1*, *Grb2*, and *Thy1*, potentially reflecting responses to the altered cell-cell interactions within the HSC niche.

We next expanded the above analysis to include distal and proximal cCREs to identify sub-cCRE age-associated alterations genome-wide. This analysis revealed 4 clusters (n = 18,166) with 84% of age-associated sub-cCREs gaining activity during aging (**Fig. 4m**), denoting a global widening of cCREs. Further categorizing distal and proximal sub-cCREs as Class I or Class II, revealed Class II elements were more proximal to promoters (*p* = 1.39*10^-9^). We observed both the significant gain in expression and downstream displacement of nucleosomes occluding motifs associated with AP-1 (Fosl2, Fos, Fosb, Jund and Junb) and Tcf4 TFs (**Extended Data Fig. S8b**), which has been reported by previous studies to be involved in HSC aging^36, 43^. Similarly, we also observed down-regulation of TFs such as Arnt and Atf7^36, 43^. Analysis of Class II elements revealed Hif1a (heat shock), Smad3 (cell adhesion), and Ybx1/3 (proteostasis) regulators. In contrast, Class I elements reflected alterations to TFs such as Hox, Rorc, Maf and Runx factors.

Overall, we find that aging is accompanied by widespread widening of cCREs to expose new TF binding sites. These sub-cCRE changes are particularly enriched at loci encoding genes involved in regulating proteostasis and cell adhesion in HSCs, and constitute a different class of regulation (Class II) than the Class I regulation that has commonly been seen in development and cell fate decisions, which is driven by opening and closing of entire cCREs (**Fig. 4n**). These data thus point to a new mechanism underlying age-dependent alterations in gene expression, and help to explain why certain HSC functions, including protein quality control, cell adhesion and RNA processing, are particularly vulnerable to age-dependent decline.

## Discussion

Our results highlight limitations of treating cCREs as digital, indivisible units. The observation that cCREs shrink, expand, and merge as cells modulate the activity of sub-cCREs argues for a model wherein cCREs dynamically recruit new effector proteins to alter their function over time. Prior studies argued that TF binding is determined by wholesale opening or closing of cCREs instead of differential binding of TFs within the same cCRE^44^. In contrast, our study describes structural changes to cCREs, mediated by the repositioning of nucleosomes and exposure of previously inaccessible DNA for TF binding. This difference likely arises from the increased resolution with which we were able to examine cCRE structure, including the ability to footprint objects of various sizes along a continuous trajectory of cell differentiation. In support of this model, studies mapping TF binding by ChIP-seq report that TFs switch in development^3, 45, 46^. Further, prior studies using high resolution ChIP-seq find that nucleosomes are in active competition with transcriptional machinery^7^.

We find that cCREs may be divided into sub-cCREs according to their change across single-cell data to significantly improve mapping of chromatin accessibility. In hematopoiesis, we find that individual cCREs change in structure, exposing DNA for TF binding, and altering gene expression. In aging, we find that most age-associated promoter changes alter the structure of cCREs, while fewer alter overall accessibility of the element. Our approach for footprinting is generalizable, and may reveal sub-cCREs in previously published bulk or single-cell ATAC-seq data, creating immediate opportunities across diverse studies of healthy and disease biology. From this vast repertoire of regulatory diversity, we anticipate discovering functions for not yet appreciated chromatin remodelers that slide or evict nucleosomes from regulatory DNA. Parsing cCREs into sub-cCREs may also ascribe new functions to disease-causing genetic variation previously overlooked by peak-based analyses. Taken together, our approach reveals the dynamics and functional importance of cCRE structure, providing new insights into gene expression and highlighting functional DNA at single-cell and single-base-pair resolution.

## Extended Data Figures

**Extended Data Figure S1.**
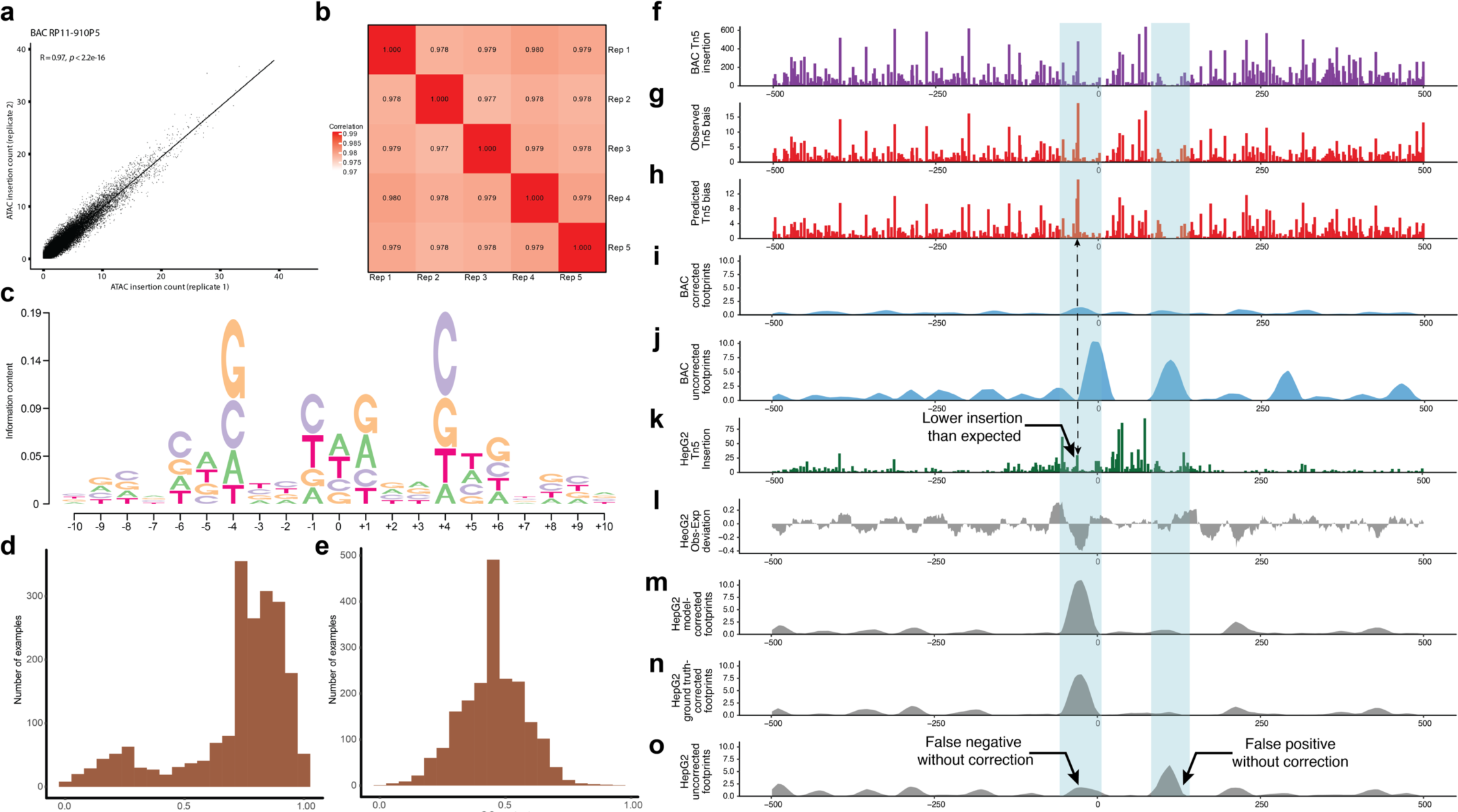
Tn5 bias modeling and footprinting. **a**, Scatter plot comparing single nucleotide observed Tn5 insertion bias on BAC RP11-910P5 from replicate 1 and 2. **b**, Heatmap showing Pearson correlation of observed Tn5 on all BACs among replicates. **c**, Motif plot of Tn5 sequence bias. **d**, Histogram of local GC-content in a +/−10 bp window for top 2000 genomic positions where the neural network Tn5 bias model achieved the highest improvement in prediction error compared to the PWM bias model. **e**, Histogram of local GC-content in a +/−10 bp window for bottom 2000 genomic positions where the neural network Tn5 bias model achieved the least improvement in prediction error compared to the PWM bias model. **f-o**, Testing our footprinting framework in an example cCRE region. **f-j**, Results for BAC naked DNA. **f**, Observed raw Tn5 insertion counts. **g**, Observed Tn5 bias. **h**, Tn5 bias predicted by the convolutional neural network. **i**, Footprint scores with bias correction. **j**, Footprint scores without bias correction. **k-o**, Results for HepG2 chromatin ATAC-seq. **k**, Observed raw Tn5 insertion counts. **l**, Observed-expected deviation of center / (center + flank) insertion ratio. **m**, Footprint scores with model-based bias correction. **n**, Footprint scores with bias correction using ground truth bias in **g**. **o**, Footprint scores without bias correction.

**Extended Data Figure S2.**
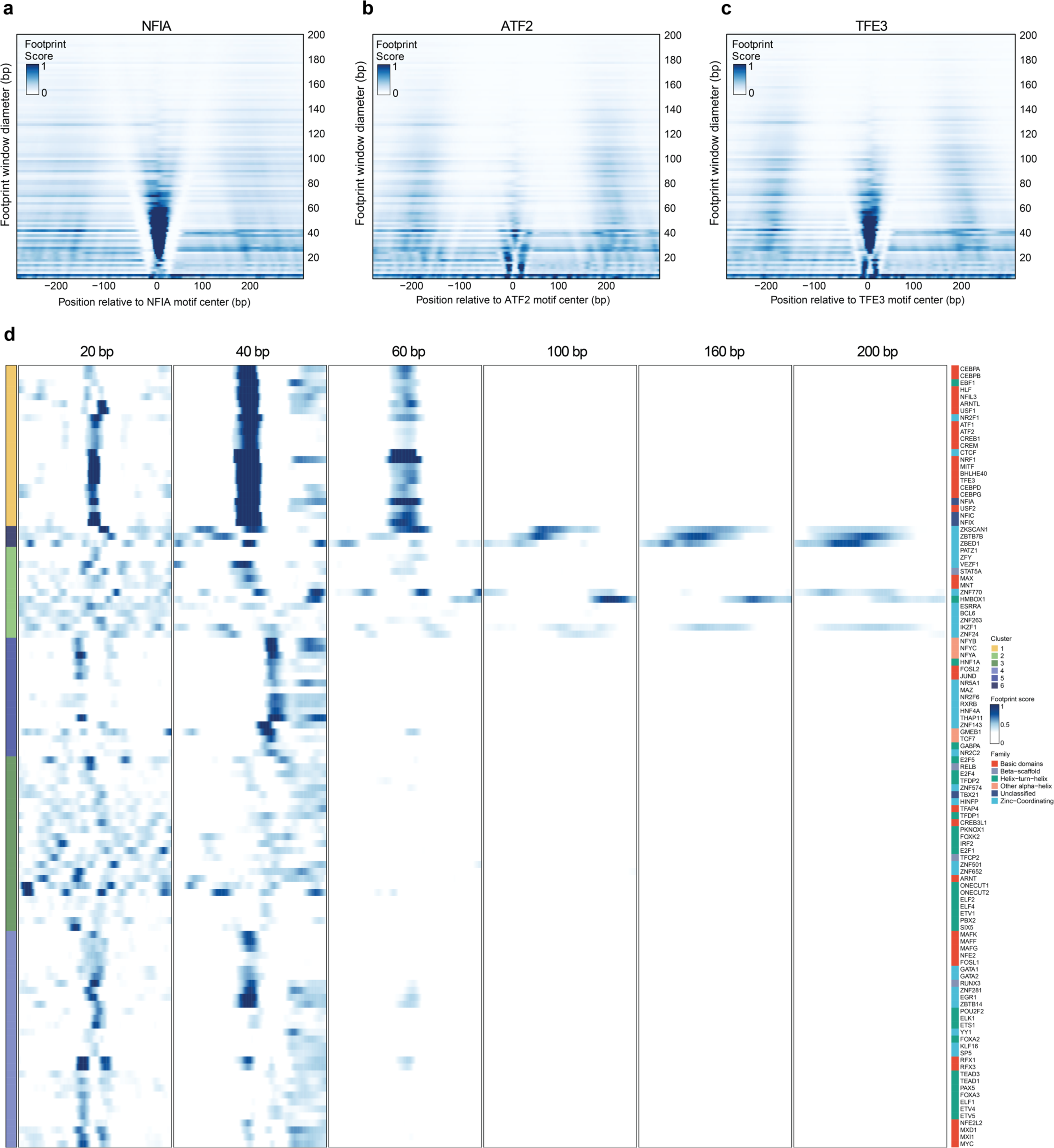
Multi-scale aggregate footprints centered around different TF motif sites. **a-c,** Multi-scale footprints for example TFs. **a,** NFIA. **b**, ATF2. **c**, TFE3. **d**, Heatmap showing clustering of multi-scale aggregate footprints of different TFs. Each row is the multi-scale aggregate footprints of a specific TF. Left color bar shows the cluster each TF is in. Right color bar shows the TF family each TF belongs to.

**Extended Data Figure S3.**
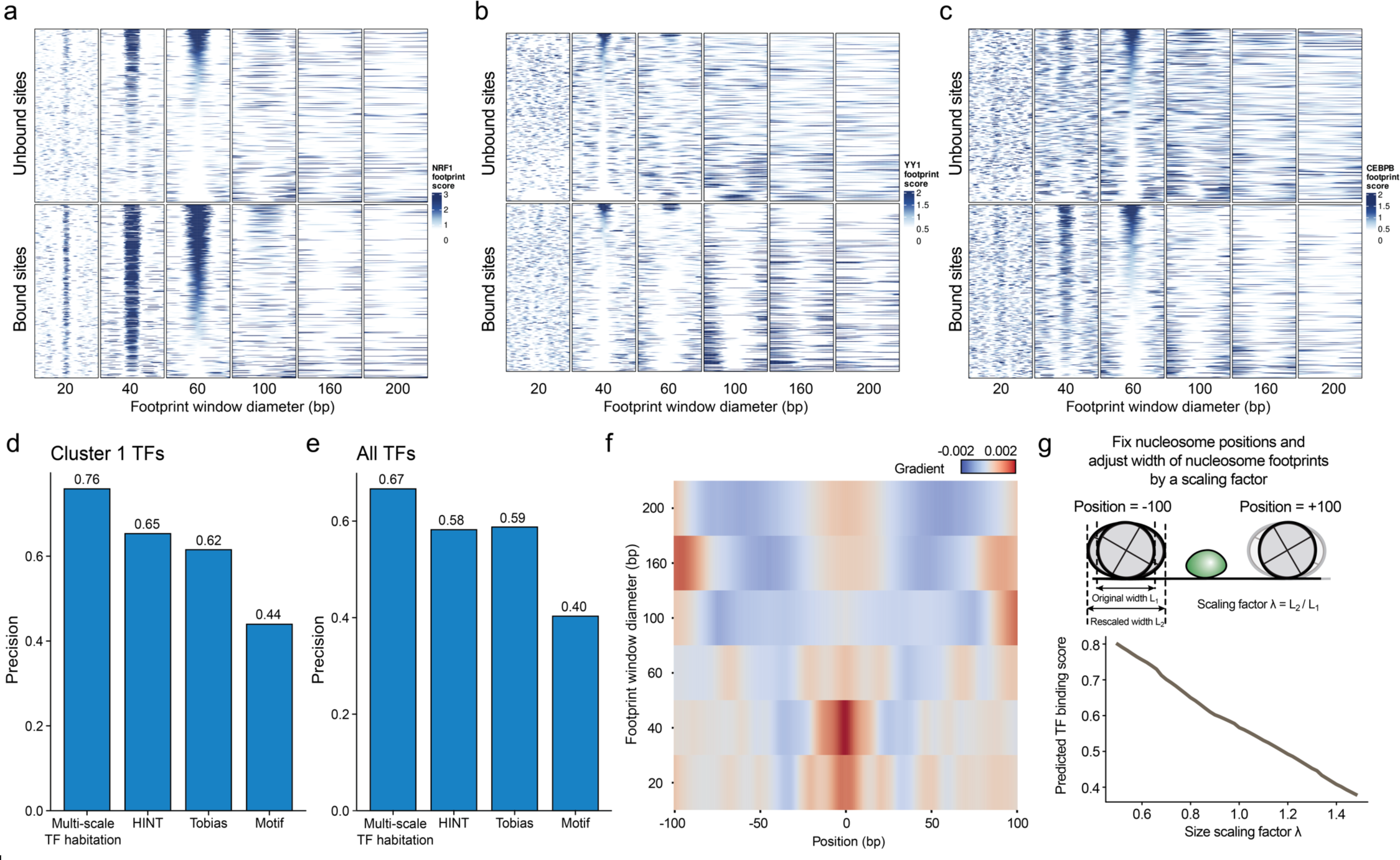
Predicting TF binding using multi-scale footprints. **a-c,** Multi-scale footprints around individual bound and unbound TF motif sites similar to Figure 2D. **a**, NRF1. **b**, YY1. **c**, CEBPB. **d**, Bar plot showing performance of different methods when benchmarked on cluster 1 TFs. **e**, Bar plot showing performance of different methods when benchmarked on TFs from all clusters. **f**, Heatmap showing gradients of predicted TF binding score with respect to input multi-scale footprints. Rows correspond to different footprint scales and columns represent single base pair positions within a +/−100 bp range from the center. **g**, Effect of changing nucleosome footprint width on predicted TF binding scores. The two nucleosomes are fixed at +/−100 bp positions, respectively and their widths are scaled by a scaling factor λ.

**Extended Data Figure S4.**
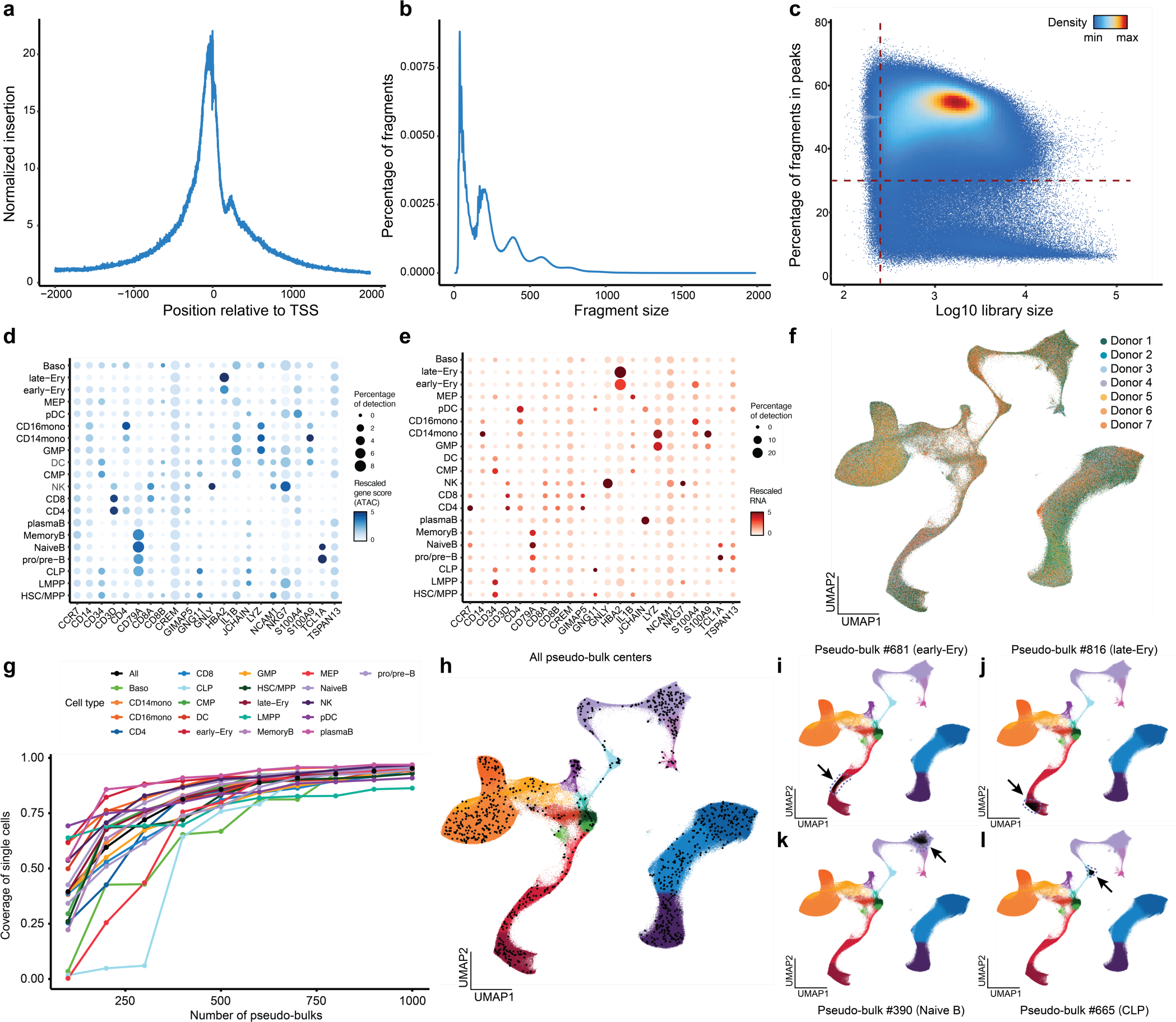
Quality control and pseudo-bulking of the human bone marrow dataset. **a**, Tn5 insertion enrichment around TSSs. **b**, Fragment size distribution. **c**, Scatter plot showing library size and fraction of reads in peaks (FRIP) of single cells. **d**, Dot plot showing gene scores (ATAC signal within a region around promoter) of marker genes across cell types. **e**, Dot plot showing of RNA levels of marker genes across cell types. **f**, UMAP showing donor origin of single cells. **g**, Line plot showing single cell coverage for each cell type as a function of the number of pseudo-bulks. **h**, UMAP showing the positions of pseudo-bulk centers for all 1000 pseudo-bulks we generated. **i-l**, Example pseudo-bulks. Black dots represent member cells within the pseudo-bulk.

**Extended Data Figure S5.**
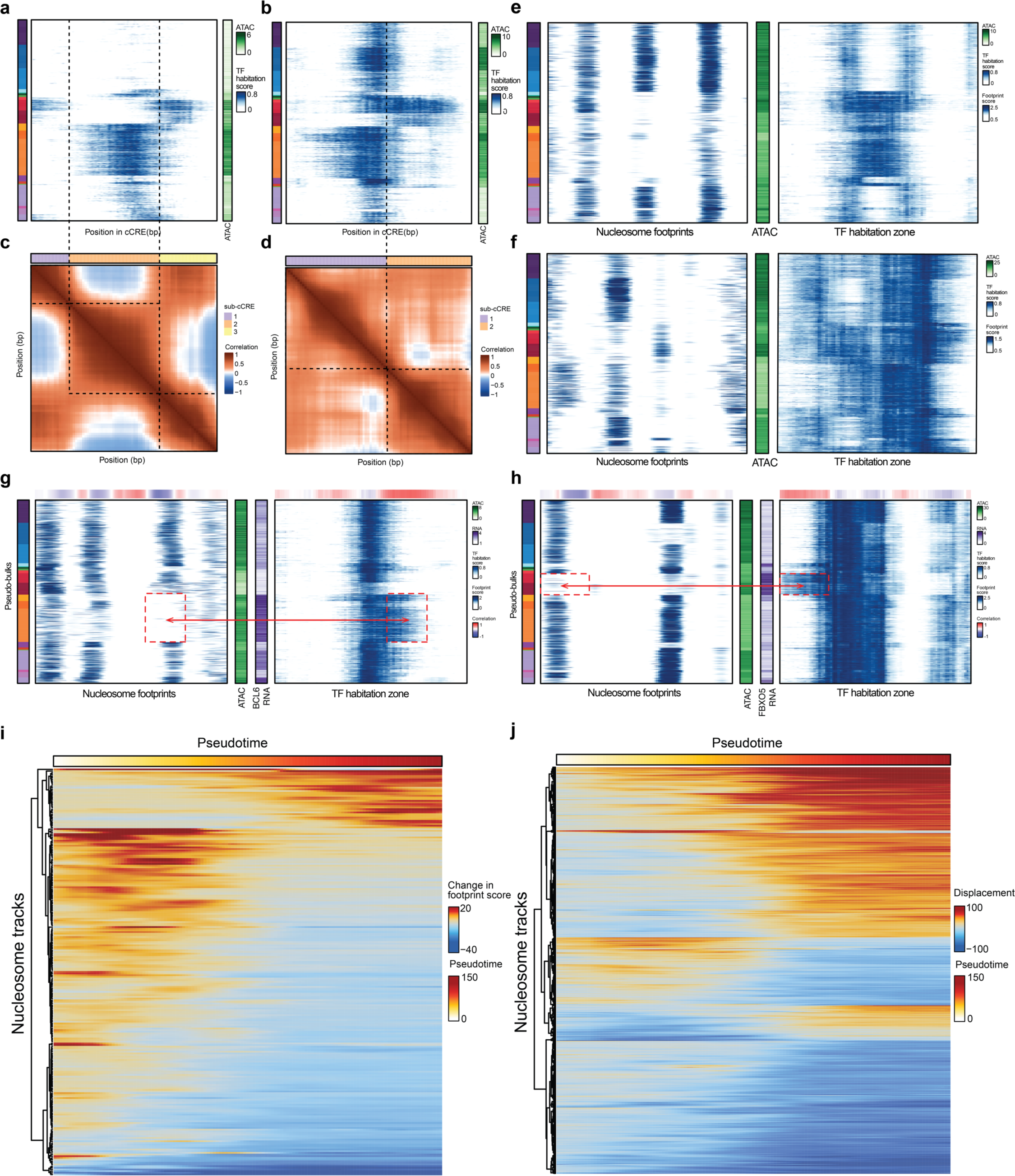
sub-cCRE and nucleosome dynamics. **a-d**, Defining sub-cCREs. **a-b**, Heatmap of predicted TF habitation scores in the cCRE at chr17:40329944-40330943 and chr20:56411666-56412665, respectively. Each row corresponds to a single pseudo-bulk, while each column represents a single base pair position in the cCRE. Left color bar shows the cell type labels of pseudo-bulks, and the colormap is the same as in Figure 3a. Right color bar shows total accessibility of the cCRE across the pseudo-bulks. **c-d**, Heatmap showing pairwise correlation of TF habitation scores among individual base pair positions within the same cCRE as in **a-b**. Top color bar shows results of sub-cCRE segmentation. **e-f**, Examples of cCRE merging and unmerging. **e**, Example cCRE at chr11:64810186-64811185. **f**, Example cCRE at chr1:84690049-84691048. Left heatmaps show the nucleosome footprints (footprint scores calculated at 100 bp scale), while right heatmaps show the TF habitation scores in the same region. Each row corresponds to a single pseudo-bulk, while each column represents a single base pair position in the cCRE. Middle color bar shows total accessibility of the cCRE across the pseudo-bulks. **g-h**, Example cCREs with strong sub-cCRE-gene correlation and weak cCRE-gene correlation. **g**, Example cCRE at region chr3:187739617-187740616. **h**, Example cCRE at region chr6:152982810-152983809. Middle color bars show total accessibility of the cCRE and the RNA of the target gene, respectively. **i-j**, Tracking nucleosome dynamics across erythroid differentiation. **i**, Heatmap showing nucleosome binding/eviction dynamics across pseudotime during erythroid differentiation. Rows are individual nucleosome tracks. Color represents the change in footprint score compared to t = 0. **j**, Heatmap showing nucleosome sliding dynamics across pseudotime during erythroid differentiation. Rows are individual nucleosome tracks. Color represents the displacement (in bp) compared to starting position at t = 0. Negative values represent sliding towards upstream regions and vice versa.

**Extended Data Figure S6.**
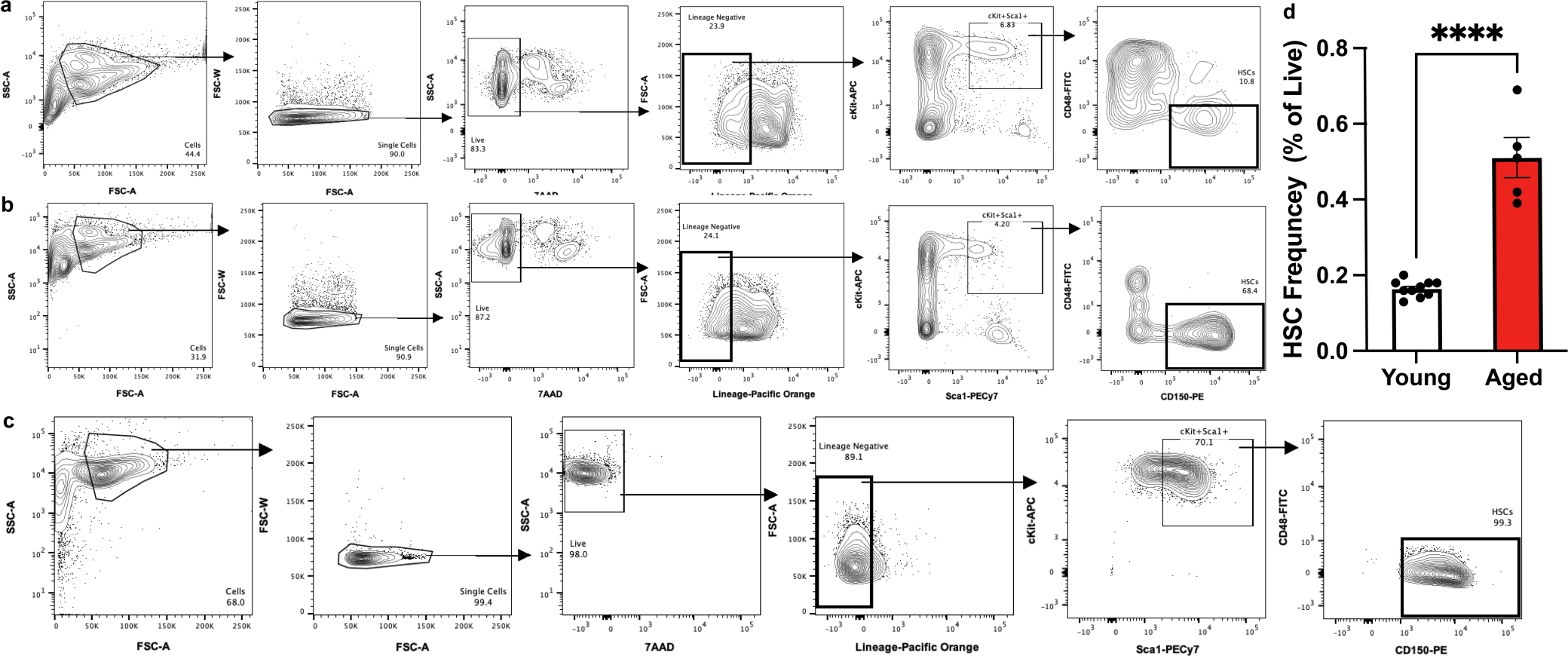
FACSorted Hematopoietic Cells from Aging Male Mice. **a-b**, Flow cytometry gating strategy for isolation of hematopoietic progenitor cells (Lineage Negative, gate bolded; Live Lin^-^) and hematopoietic stem cells (HSCs, gate bolded; Live Lin^-^ Sca1^+^ cKit^+^ CD48^-^ CD150^+^) from the bone marrow (BM) of young (**a**, n = 10) and aged (**b**, n = 5) male C57BL6/J mice. Representative FACS plots shown from one young and one aged mouse. For individual FACS plots from each mouse, see Extended Data File 1. **c**, Purity of resorted HSCs was greater than 99%. **d**, Frequency of FACSorted HSCs in young and aged mice (two-tailed t-test; t13 = 9.283, p<0.0001).

**Extended Data Figure S7.**
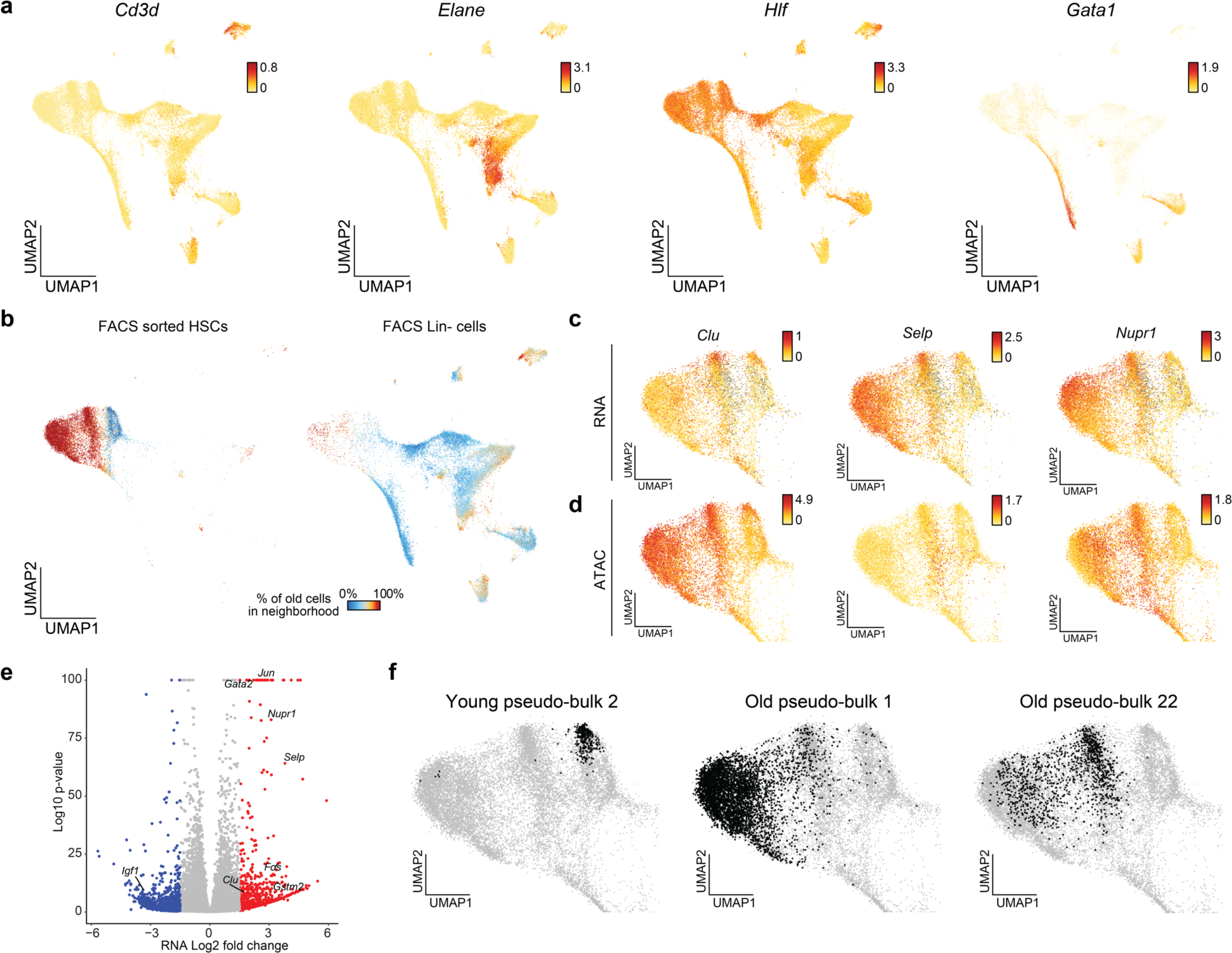
Characterizing age-related changes in mouse HSCs. **a**, UMAP showing gene scores of cell type marker genes *Cd3d*, *Elane*, *Hlf*, and *Gata1*, respectively. **b**, Percentage of old cells in the 100-cell nearest neighborhood for FACS sorted HSCs (left) or Lineage-cells (right). **c-d**, UMAP showing (**c**) RNA and (**d**) ATAC levels of aging marker genes (*Clu*, *Selp*, *Nupr1*) in HSCs. **e**, Volcano plot of differential RNA testing (old-versus-young). **f**, UMAP showing example pseudo-bulks. Black dots represent member cells within each pseudo-bulk.

**Extended Data Figure S8.**
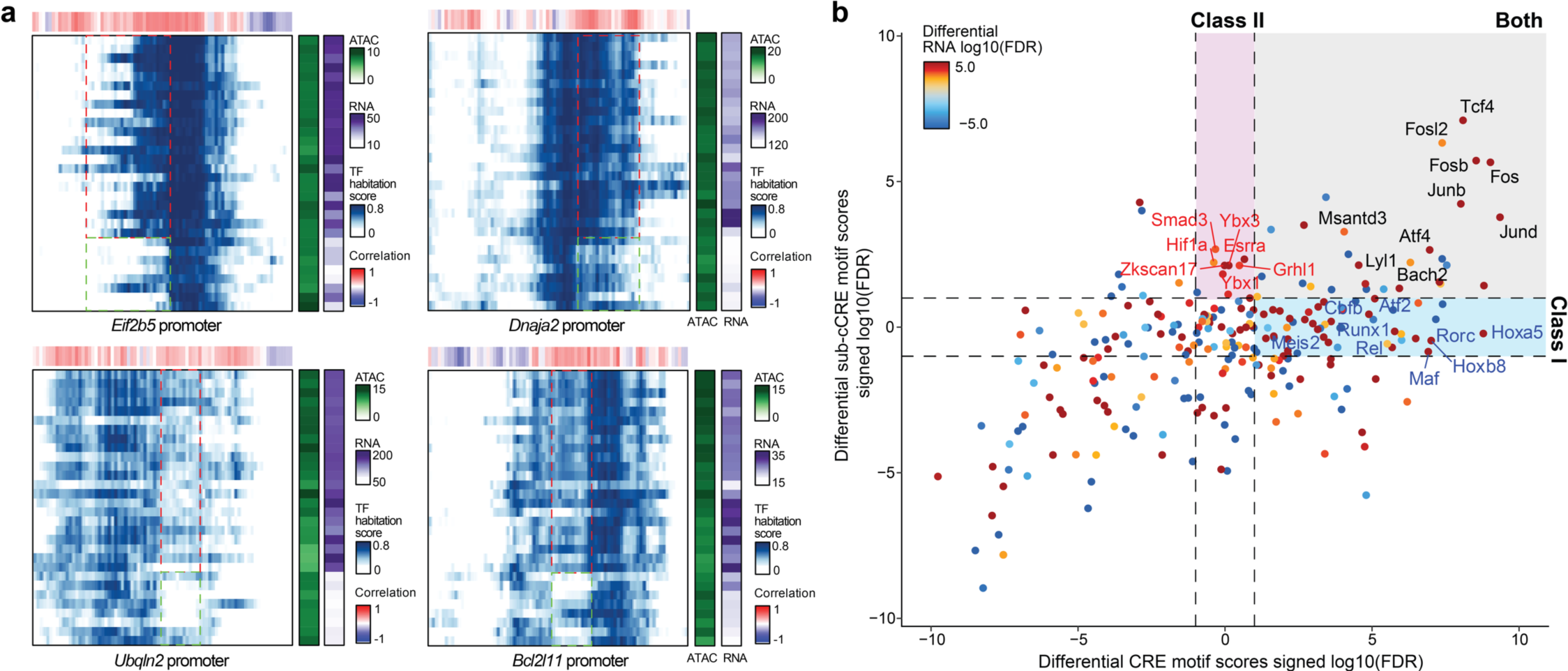
Hallmarks of aging-associated sub-cCRE alterations. **a**, Heatmaps of predicted TF habitation scores in the promoters of *Eif2b5*, *Dnaja2*, *Ubqln2*, and *Bcl2l11*, respectively. Each row corresponds to a single pseudo-bulk, while each column represents a single base-pair position in the cCRE. Right color bar shows total accessibility of the cCRE and RNA level of the corresponding gene across the pseudo-bulks. Top color bar shows correlation of TF habitation score with RNA level at each base-pair position. **b**, Scatter plot comparing differential TF motif score testing results using cCREs and sub-cCREs as input features, respectively.

## Methods

### EXPERIMENTAL METHODS

#### Cell culture

HepG2 cells were cultured in Dulbecco’s Modified Eagle Medium (DMEM, 11965092, ThermoFisher) with the addition of 10% FBS and 1% of penicillin-streptomycin. Cells were incubated at 37°C in 5% CO_2_ and maintained at the exponential phase. Cells were digested with TrypLE express (12604013, ThermoFisher) for preparing single-cell suspension.

V6.5 mouse embryonic stem cells were cultured in Glasgow Minimum Essential Medium (GMEM) supplemented with 10% FBS, 2 mM L-glutamine, 1% Pen/Strep, 1 mM sodium pyruvate, 2000 units/mL (10ng/mL) Leukemia Inhibitory Factor (LIF, Millipore), 1x Minimum Essential Medium Non-Essential Amino Acids (MEM NEAA, Invitrogen) and 50 uM β-Mercaptoethanol. Tissue culture plates were coated with 0.2% gelatin and 0.25 mg/mL laminin for 2 hours at 37C before seeding. Media was changed every other day, and cells were split every 3-4 days.

#### BMMC sample processing

Frozen human Bone Marrow Mononuclear Cells (BMMCs, Allcells) were thawed in a 37 °C water bath for 1 min and transferred to a 15 mL centrifuge tube. 10 mL of pre-warmed DMEM with 10% FBS was added to cells drop-wisely. The cells were spun at 400g for 3 min at room temperature. After removing supernatant, the cells were washed twice in 0.5 mL PBS with 0.04% BSA. To deplete neutrophils, the cells were resuspended in 100 μl chilled DPBS with 0.2% BSA and 10 μl of human TrueStain FcX (BioLegend, 422302) and incubated on ice for 10 min to reduce non-specific labeling. The cells were then incubated on ice for another 30 min after adding 0.5 μl of biotin anti-human CD15 antibody (BioLegend, 301913). After immunostaining, 25 μl of MyOne T1 beads were added to the sample to capture the neutrophils for 5 min at room temperature. We then added 900 μl of DPBS with 0.2% BSA to dilute the sample. The sample was placed on a magnet for 3 min and 1 ml of the sample was transferred to a new tube while the sample was on the magnet. The cells were ready for fixation and SHARE-seq experiment.

#### Fixation

Cells were centrifuged at 300g for 5 minutes and resuspended to 1 million cells/ml in PBSI. Cells were fixed by adding formaldehyde (28906, ThermoFisher) to a final concentration of 1% and incubated at room temperature for 5 minutes. The fixation was stopped by adding 56.1 μl of 2.5M glycine, 50 μl of 1M Tris-HCl pH 8.0, and 13.3 μl of 7.5% BSA on ice. The sample was incubated at room temperature for 5 minutes and then centrifuged at 500g for 5 minutes to remove supernatant. All centrifugations were performed on a swing bucket centrifuge. The cell pellet was washed twice with 1ml of PBSI, and centrifuged at 500g for 5 minutes between washings. The cells were resuspended in PBS with 0.1U/μl Enzymatics RNase Inhibitor and aliquoted for transposition.

#### SHARE-seq

Following fixation SHARE-seq was performed as previously described^26^, with the following modifications. To improve transposition, transposition was performed using pre-assembled Tn5 (seqWell, Tagify(TM) SHARE-seq Reagent). To improve RNA capture, we added polyA to transcripts prior to reverse transcription. To do this, transposed cells (60 μl) were mixed 240 μl of poly(A) mix (final concentration of 1× Maxima RT buffer, 0.25 U/μl Enzymatics RNase Inhibitor, 0.25 U/μl SUPERase RI, 0.018 U/μl E.coli poly-A enzyme (M0276L), 1 mM rATP). The sample was aliquoted to 50 μl per PCR tube and incubated at 37 °C for 15 minutes.

#### Quantification and sequencing

Both scATAC-seq and scRNA-seq libraries were quantified with the KAPA Library Quantification Kit and pooled for sequencing. Single cell libraries were sequenced on the Nova-seq platform (Illumina) using a 200-cycle kit (Read 1: 50 cycles, Index 1: 99 cycles, Index 2: 8 cycles, Read 2: 50 cycles). Bulk libraries were sequenced on the Nova-seq platform (Illumina) using a 100-cycle kit (Read 1: 50 cycles, Index 1: 8 cycles, Index 2: 8 cycles, Read 2: 50 cycles).

#### SHARE-seq data pre-processing

SHARE-seq data were processed using the SHARE-seqV2 alignment pipeline (https://github.com/masai1116/SHARE-seq-alignmentV2/) and aligned to hg38. Open chromatin region peaks were called on individual samples using MACS2 peak caller^47^ with the following parameters:--nomodel – nolambda –keep-dup-call-summits. Peaks from all samples were merged and peaks overlapping with ENCODE blacklisted regions (https://sites.google.com/site/anshulkundaje/projects/blacklists) were filtered out. Peak summits were extended by 150 bp on each side and defined as accessible regions (for footprinting analyses, thes peaks were later resized to 1000 bp in width). Peaks were annotated to genes using Homer ^48^. The fragment counts in peaks and TF scores were calculated using chromVAR^49^. Cell barcodes with less than 30% reads in peaks (FRiP) or 250 unique fragments were removed. The aligned reads were then intersected with peak window regions, producing a matrix of chromatin accessibility counts in peaks (rows) by cells (columns). To examine the cell identity, the cisTopic (50 topics)^50^ were used for dimension reduction, followed by Louvain clustering. The progenitor populations were sub-clustered to obtain finer cell identity. The data were projected into 2D space by UMAP^51^. Seurat V3^52^ was used to scale the DGE matrix by total UMI counts, multiplied by the mean number of transcripts, and values were log transformed.

#### Generation of BAC naked DNA data

We selected 25 chromatin regions based on overlap with a manually selected set of key transcription factors and differentiation related genes. The BAC clones (BACPAC Resources) were cultured in LB for 14 hours. The BAC DNA was extracted using ZR BAC DNA Miniprep Kit (Zymo, D4048) following manufacturer’s instructions. The purified DNA was quantified using Qibit (ThermoFisher). The BAC DNA were tagmented similar to the SHARE-seq ATAC-seq experiment. Briefly, 50 ng of BAC DNA from multiple clones were pooled for tagmentation following the SHARE-seq transposition condition. The tagmented DNA was purified using a Qiagen Minelute PCR clean up kit and then amplified for 7 cycles by PCR. To minimize batch effect, we generated 5 replicates and pooled all the materials for sequencing. The library was sequenced on a Nova platform (Illumina) using a 100-cycle kit (Read 1: 50 cycles, Index 1: 8 cycles, Index 2: 8 cycles, Read 2: 50 cycles). The sequencing data was processed the same way as SHARE-seq ATAC-seq data.

#### Aging Multi-ome experiment

Mouse experiments were approved and performed in compliance with Harvard University’s Institutional Animal Care and Use Committee. C57BL6 mice were obtained from either Jackson Laboratory or the National Institute on Aging Aged Rodent Colony (Charles River Laboratory), and housed at a density of 2-5 mice per cage in standard ventilated racks and provided food and water *ad libitum* in a pathogen-specific free facility accredited by the Association and Accreditation of Laboratory Animal Committee (AALAC). Mouse cages contained Anderson’s Bed o Cob bedding (The Anderson, Inc), two nestlets (Ancare, 2×2” compressed cotton square), and a red mouse hut (Bioserv). For HSC isolation and flow cytometry. Cells from the bone marrow of long bones (2 femurs and 2 tibias per mouse) from young (n = 10; 11 weeks old) and aged (n = 5; 24 mo. old) male C57BL/6 mice were flushed with a 21-gauge needle into staining media (HBSS/2% fetal bovine serum), pelleted, and resuspended in ACK lysis buffer for 5 min on ice. Cells were then washed with staining media, filtered through a 40mm cell strainer, pelleted, and incubated with the following cocktail of rat anti-mouse, biotin conjugated lineage antibodies on ice for 30 min: CD3 clone C145-2c11 (Biolegend, 1000304; 1:100), CD4 clone GK15 (Biolegend, 1000404; 1:400), CD5 clone 53-7.3 (eBioscience, 13-0051-85; 1:400), CD8 clone 53-6.7 (Biolegend, 100704; 1:400), CD19 clone 6D5 (Biolgend, 115504; 1:400), B220 clone RA3-6B2 (Biolegend, 103204; 1:200), GR1 (Ly6-G/Ly6-C) clone RB6-8C5 (eBioscience, 13-5931-82; 1:400), Mac1/CD11b clone M1/70 (Biolegend, 101204; 1:800), and Terr119 clone TERR-119 (Biolegend, 116204; 1:100). Cells were then washed in staining media, with a small aliquot reserved for each sample to serve as a non-depleted control, and lineage depleted using sheep anti-rat Dynabeads (Invitrogen, 1135) on a magnet. Cells were washed, pelleted, and incubated with the following cocktail of anti-mouse antibodies on ice for 45 min. to identify hematopoietic stem cells (HSC): Pacific Orange Streptavidin (Invitrogen, S32365; 1:500), PE/Cy7 Sca1(Ly-6a/E) clone D7 (eBioscience, 25-5981-82; 1:200), APC cKit clone 2B8 (BD Pharmingen, 553356; 1:200), FITC CD48 clone HM48-1 (Biolegend, 103403; 1:200), and PE CD150 clone Tc15-12F12.2 (Biolegend, 115904; 1:200). Following incubation, cells were washed and resuspended in staining media, and 7-AAD (BD Pharmingen, 559925; 1:50) added immediately prior to flow cytometry. Cell sorting of HSCs (Live Lin^-^ Sca1^+^ cKit^+^ CD48^-^ CD150+) was performed on a BD FACS Aria II, and data analysis performed using BD FACS Diva and FlowJo software. Data processing was performed using CellRanger.

After sorting, nuclei were isolated following 10x Genomics’ demonstrated protocol "Low Cell Input Nuclei Isolation", which is described in the CG000365 User Guide. Nuclei were then processed using the Chromium Single Cell Multiome ATAC + Gene Expression kit (10x Genomics), following manufacturer’s instructions, to obtain between 2,000 and 10,000 cells per sample. Libraries were sequenced on an Illumina Nextseq system using the following sequencing formats: Read 1 - 28, i7 index - 10, i5 index - 10, Read 2 - 44 (scRNA-seq), Read 1 - 30, i7 index - 8, i5 index - 24, Read 2 - 30 (scATAC-seq). Data processing was performed using the CellRanger software from 10x Genomics.

#### Tn5 sequence bias modeling

##### Getting Tn5 insertion counts

The ends of the fragments files are shifted by +4/-4 (in 1-based indexing system) to obtain the center of the 9 bp staggered end created by Tn5 transposition. The number of insertions at each single base-pair position within each cCRE from each sample is then quantified and stored in a sample-by-cCRE-by-position 3D tensor for fast data retrieval.

##### Data preprocessing

The model takes local DNA sequence context as input and predicts single-base pair resolution Tn5 bias. To this end, the +/−50 bp DNA sequence surrounding each position of interest is encoded by one-hot encoding into a 101-by-4 matrix and used as model input. For the prediction target, we use local relative Tn5 bias as the target value. More specifically. The raw Tn5 insertion count at each position is divided by the average Tn5 insertion count within a +/-50 bp window. Positions with low local coverage (< 20 insertions per bp) were removed to guarantee quality of training data. To facilitate model training, the resulting observed Tn5 bias values are log10-transformed and rescaled. For dataset partition, we randomly split all the BACs into 80%, 10%, and 10% for training, validation, and test sets. In other words, all data originating from the same BAC belong to the same partition. This is to prevent overlapping local sequence contexts ending up in both training and testing datasets, which might lead to overestimation of performance. To guarantee equal coverage of examples with different bias levels, we binned all training examples into 5 bins based on their Tn5 bias values, and up-sampled each bin so that all bins end up with the same number of examples. Additionally, given the symmetric nature of Tn5 insertion, we generated reverse complement sequences of the training examples as data augmentation. The original and reverse complement data were combined, shuffled, and then used for model training.

##### Model architecture

The convolutional network consists of three convolution & max-pooling layers and two fully connected layers. Each convolution and max-pooling layer performs convolution, ReLU nonlinear activation^53^, and max pooling sequentially. We used 32 filters of width 5 for each layer, along with “same” padding mode and stride size of 1. The two following fully connected layers have output dimensions of 32 and 1, respectively. ReLU activation is used by the first fully connected layer and linear activation is used by the second layer (i.e., the final output layer).

##### Model training and evaluation

The model was trained on the training set, and hyperparameters were optimized based on performance on the validation set. Final performance of the frozen model was evaluated on the test set. The model was implemented using Keras^54^, trained with mean square error as loss function and optimized using the Adam optimizer^55^ with default parameters. Training was performed with a batch size of 64 and early stopping based on model loss on the validation set.

##### Benchmarking with other Tn5 bias models

Methods including k-mer models (k = 3, 5, 7) and PWM methods (single nucleotide and dinucleotide) were included in benchmarking. For k-mer methods, the foreground and background frequencies for all possible k-mer sequences were quantified. The foreground frequency / background frequency ratio was used as the estimated Tn5 bias for the corresponding k-mer. For single nucleotide PWM, we calculated foreground and background base frequencies within a +/−10 bp window (total length = 21) and computed the PWM of Tn5 insertion. Dinucleotide PWM scores were calculated using TOBIAS^21^ with default settings.

##### Genome-wide Tn5 bias reference tracks

Sequences of reference genomes for *Homo sapiens* (hg38), *Mus musculus* (mm10), *Drosophila melanogaster* (dm6), *Saccharomyces cerevisiae* (sacCer3), *Caenorhabditis elegans* (ce11), *Danio rerio* (danRer11), and *Pan troglodytes* (panTro6) are downloaded from the UCSC genome browser website^56^ https://hgdownload.soe.ucsc.edu/goldenPath/. The aforementioned Tn5 bias neural network model was applied to each position in the reference genomes to generate genome-wide Tn5 bias tracks.

#### Computing footprint scores

To detect DNA-protein interactions at different scales within cCREs, we implemented a framework for computing footprint scores for each base pair position in the cCRE. In short, for each single bp position, we define a center footprint window and flanking windows on both sides (**Figure 1e**). Then we calculate the observed ratio of center / (center + flanking) Tn5 insertion counts. The foreground observed ratio is compared to a background distribution to calculate statistical significance, which is then converted to a footprint score.

##### Estimation of background dispersion

Given a specific combination of center bias, flanking bias, and local coverage, we expect a certain distribution of center / (center + flanking) insertion ratio when no protein is bound. This is defined as the background distribution. Such background distribution can be estimated using BAC naked DNA Tn5 insertion data. To this end, we first randomly sampled 100,000 positions from the BAC dataset, and retrieved their local coverage (defined as the total insertion number in center and flanking areas), center bias, as well as flanking bias. Then for each sampled position A, we identified 500 nearest neighbor positions NN_1_-NN_500_ in the 3-dimensional space of (center bias, flanking bias, local coverage). To make sure each dimension is weighed equally, the values of each dimension were first normalized to zero mean and unit variance. The 500 nearest neighbor observations can be considered as background observations with nearly identical bias and coverage, and the center / (center + flanking) ratio of NN_1_-NN_500_ forms the background distribution of position A. Therefore, for each of the 100,000 sampled positions, we can calculate the mean and standard deviation of its background ratio distribution. This allows us to train a background dispersion model that takes the tuple (center bias, flanking bias, local coverage) as input and predicts the mean and standard deviation of the background distribution very efficiently. To make sure the model is exposed to training examples with a wide range of local coverage, we down-sampled the BAC dataset to 50%, 20%, 10%, 5%, and 1% of the original sequencing depth. Finally, we trained a neural network with a single hidden layer (32 nodes, ReLU activation^53^) and linear output layer activation. The dataset was randomly split into 80% training, 10% validation, and 10% test. The model was implemented using Keras^54^, and trained on the training dataset with mean squared error loss using the Adam optimizer^54, 55^. Early stopping was determined using loss on the validation set, and performance of the final model was evaluated on the test set. Additionally, we trained separate models for each footprint radius due to the drastic differences in total center or flank bias when footprint radius varies. For details, see Supplementary Notes.

##### Calculating footprint scores

For each position in the cCRE, we define a center footprint window and flanking windows on both sides. We first calculate the foreground observed center / (center + flanking) ratio of Tn5 insertion counts. Then we apply the pre-trained background dispersion model to calculate the mean and standard deviation of its background distribution. We next use a lower-tailed *z*-test to calculate the *p*-value for footprinting. If the observed ratio is significantly lower than the background distribution, then this position is likely to be bound by a protein. More specifically, to avoid calling footprints at positions where only one flanking side shows higher Tn5 insertion than the center window but not the other, we perform center-versus-left and center-versus-right tests separately and keep the larger *p*-value (See Supplementary Notes for details). The-log10(*p*-values) are smoothed by running-max and running-mean smoothing and then used as the final footprint scores.

##### Aggregate footprinting

To calculate aggregate footprints, Tn5 insertions surrounding TF or nucleosome binding sites across the genome are first aggregated and then used to calculate footprint scores. For TFs, we selected sites with a matched TF motif using motifmatchr^57^ (p.cutoff = 1e-5) and overlapping with a ChIP-seq peak of the corresponding TF. For motif matches on the minus strand, the Tn5 insertion profile surrounding the motif is inverted so the insertions for different sites are aligned in the same direction. For nucleosomes, we downloaded a previously published list of chemically mapped nucleosome positions in mouse embryonic stem cells (mESCs)^24^ and used these positions for aggregating nucleosome footprinting with mESC single cell ATAC-seq data.

#### Predicting TF binding

##### Input data

To predict the landscape of TF binding, we trained a binary classifier that predicts whether any TF motif site is bound by the corresponding TF. Motif sites are identified by the matchMotifs function in the motifmatchr package^57^. All sites with a matching *p*-value below 5e-5 are kept. For any TF motif site, we use multi-scale (20 bp, 40 bp, 60 bp, 100 bp, 160 bp, 200 bp in diameter) footprints within a +/−100 bp local area centered around the motif, as well as a motif match score as input to the model. The motif match score returned by the matchMotifs function is quantile-transformed to uniform distribution. As a result, by combining the 201-dimensional footprint vectors from 6 different scales with a single motif match score, we end up with a 1207-dimensional vector as the final model input. The first 1206 dimensions of footprint scores are standardized individually to zero mean and unit variance. For the prediction target, we assign a label of 1 to all sites overlapping with a ChIP peak of the same TF, and a label of 0 to sites not overlapping with ChIP.

In total, we trained two separate models. (1) The first model was trained using only data of cluster 1 TFs. This model was trained to predict binding of TFs that leave strong footprints. Some TFs were found to have a very low percentage of motif sites overlapping with ChIP (< 25%), potentially due to low quality of the motif or the ChIP dataset. Such TFs are removed from model training and testing. We also added an equal number of random negative examples as well as reverse-complement examples for data augmentation. (2) The second model, referred to as the TF habitation model, was trained on TFs from all clusters (to include more training data, we were keeping TFs with > 20% of motifs overlapping with ChIP data). The model was trained to infer binding for both strong-footprinting and weak-footprinting TFs. Similarly, we added reverse complement examples for data augmentation.

For data partition, we used HepG2 SHARE-seq data and GM12878 SHARE-seq data (GM12878 data is previously published in the original SHARE-seq paper^26^) for model training and validation, and test the model on K562 Biorad single cell ATAC data, as well as three cell types (naive B cells, CD14 monocytes, and late-erythroid cells) in the human BMMC SHARE-seq dataset. In particular, for the cluster I-specific model, TFs in cluster I were used as training data and other TFs were used as validation. For the TF habitation model, HepG2 data was used as training data and GM12878 data was used as validation. After fixing model hyperparameters, HepG2 and GM12878 data were combined to train a final TF habitation model for performance testing.

##### Model architecture and training

The TF binding prediction model is a neural network model with two hidden layers (32 + 16 nodes for cluster-I specific model, and 128 + 16 nodes for the TF habitation model). ReLU activation^53^ is used by both hidden layers and sigmoid activation is used by the final output layer. The model was implemented using Keras^54^. The model was trained on the training dataset with a batch size of 128 using the Adam optimizer^58^. Binary cross entropy is used as the loss function. Early stopping was used based on model loss on the validation set.

##### ChIP validation and benchmarking with previous methods

To evaluate model performance, we used ChIP-seq as ground truth and validated predicted binding events. HepG2 and GM12878 data for model training were downloaded from ENCODE^6^. ChIP-seq for BMMC cell types were downloaded from cistromeDB^59^. For benchmarking with previous methods, to make sure we only include high quality TF binding sites, we downloaded K562 ChIP-based TF binding data from unibind^25^ (https://unibind.uio.no/search). For ENCODE datasets, we removed those with the two most severe levels of audit categories. For cistromeDB datasets, we applied QC filters as specified on the cistromeDB website http://cistrome.org/db/#/about. More specifically, we included the below filters: FRiP >= 0.01, FastQC >= 0.25, uniquely mapped ratio >= 0.6, peaks with fold change above 10 >= 500, peaks union DHS ratio >= 0.7, and PBC >= 0.8. Datasets with the below cell type labels are included: “Monocyte”, “B Lymphocyte”, “Erythroid cell”, “Erythroid Progenitor Cell”, and “Erythroid progenitor”.

The K562 datasets from unibind were used for benchmarking with previous methods, including HINT-ATAC and TOBIAS. In short, the same ATAC-seq data was used as input to all three methods. To guarantee fair comparison, we first took the intersection of candidate TF binding sites from all three methods. Then for each method, we ranked the remaining candidate sites by predicted binding score, and evaluated precision of prediction using the top 10% sites. Only TFs with at least 10% of motifs overlapping with unibind validated TF binding sites were included. Visualization of predicted and ground truth binding sites was done with the Gviz package^60^. Furthermore, to evaluate the false positive rate of each model, we also tested all three models on our BAC naked DNA data. The same data was used as input to each model and the number of predicted binding events are used to represent the false positive predictions.

##### Model Interpretation

To interpret how the TF binding model makes predictions, we first calculated gradients of the output with respect to input features using GradientTape from tensorflow^61^. Gradients were computed for each individual motif site and then the gradients for all sites were averaged and smoothed to generate the final gradient map. Additionally, we performed ablation tests to evaluate the contribution of TF and nucleosome footprints. Here, we masked TF footprints (20 bp, 40 bp, and 60 bp footprints) or nucleosome footprints (100 bp, 160 bp, and 200 bp footprints) separately during training by setting the corresponding features to zeros. The performances of the models trained on masked data were then compared to the model trained on unmasked data. Furthermore, we performed simulation analysis to study the impact of nucleosome positioning and width on model prediction. To simulate TF and nucleosome footprints that reflect their real sizes, we first generated a gaussian signal to represent the initial guess for their size (30 bp for TFs, 50 bp for nucleosomes), Next, we went through footprints called on the HepG2 dataset at the corresponding scale and found examples that highly correlates (Pearson correlation > 0.8) with our initial guess profile. The matched data was eventually averaged to get the realistic width of footprints at each scale. We next simulated a TF signal at the center of the motif, as well as signals of two flanking nucleosomes. By changing either the positioning or width of the flanking nucleosomes, we were able to evaluate their impact on TF binding by observing the changes in the predicted TF habitation scores.

#### Segmentation of sub-cCREs

cCREs are segmented into sub-cCREs using an approach conceptually similar to segmentation algorithms for topologically associating domains (TADs)^62^. In brief, each cCRE is first divided into 10 bp intervals as candidate binding sites. Suppose there are *k* intervals in a specific cCRE. We first compute TF habitation scores for these *k* candidate binding sites across all *n* pseudo-bulks, resulting in a *k*-by-*n* matrix *S* of TF binding scores. We then calculate the pairwise correlation among the rows, obtaining a *k*-by-*k* correlation matrix *M*. Then for each TF site, we assign a score to it indicating whether it should be a sub-cCRE boundary. More specifically, for each TF site, we first calculate 3 separate scores representing the average correlation (1) within the upstream neighboring TF sites, (2) across the current TF site, and (3) within the downstream TF sites. With a local neighborhood of radius r, we have:

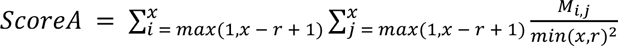

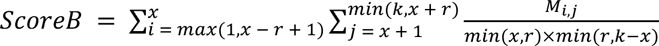

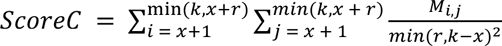

The boundary score is defined as

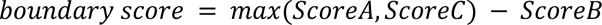

We then smooth the boundary score and identify local maximum positions as the final sub-cCRE boundaries. Furthermore, the average predicted TF habitation scores within a sub-cCRE is used as the activity of the latter. For details on cCRE segmentation, see Supplementary Notes.

#### Calculating cCRE-gene correlation and sub-cCRE-gene correlation

To investigate whether sub-cCREs can be associated with gene expression independent from the cCRE they reside in, we compared cCRE-gene correlation and sub-cCRE-gene correlation. For each gene, we identified cCREs within a 50 kb radius from the gene TSS. We then calculated correlation between cCRE accessibility (as quantified by ATAC signal within the 1 kb window) and RNA level of the gene across pseudo-bulks. Meanwhile, we also took sub-cCREs detected within the same cCRE and correlated their activities with RNA of the same gene. Low signal sub-cCREs with an activity < 0.3 were removed. To also assign statistical significance to each cCRE-gene pair or sub-cCRE-gene pair that we correlated, we constructed a background distribution of correlation values. In the case of cCREs, we first randomly selected 100,000 cCRE-gene pairs and calculated their observed correlation. For each cCRE-gene pair, we also recorded average gene expression across pseudo-bulks, average cCRE accessibility across pseudo-bulks, and GC content of the cCRE as the 3 main features of the cCRE-gene pair. Therefore, each pair will have a unique coordinate in the 3-dimensional feature space (i.e., accessibility, expression, GC content). Next, for each cCRE-gene pair that we wish to assign significance to, we find its 100 nearest neighbors in the 3-dimensional feature space. The correlation values of the nearest neighbor cCRE-gene pairs are used as the background distribution. A *z*-test is performed to obtain a *p*-value for the pair of interest. In the case of sub-cCREs, a similar procedure is conducted. The only difference is that we use mean activity of the sub-cCRE instead of mean accessibility as a main feature. Again for every sub-cCRE-gene pair 100 background pairs are found in the 3-dimensional feature space and a *z*-test is performed to get the p-value.

#### Tracking TF binding dynamics across human hematopoiesis

##### Generation of pseudo-bulks

Single cells in the human BMMC dataset were first embedded into lower dimensional space using cisTopic^50^, and then grouped into 1000 pseudo-bulks based on their spatial proximity in the cisTopic space. More specifically, we first sample 1000 cells as pseudo-bulk centers, and then identify k-nearest neighbors (k = 5000) of each center cell in the cisTopic space as other members of the same pseudo-bulk. We reasoned that sampling center cells with low local connectivity can help increase coverage of the state space by preventing over-sampling of densely connected local neighborhoods. Therefore, we first randomly sampled 10,000 scaffold cells and used them to construct a KNN graph (k = 10). Then we selected the 1000 cells with the lowest in-degree in the KNN graph as pseudo-bulk centers.

##### Computing pseudo-time

Pseudo-time along human hematopoietic lineages was computed using the Palantir package^63^. To reduce computing time, we randomly sampled 100,000 cells from the human BMMC dataset as scaffold cells. The cisTopic embedding of the scaffold cells as well as pseudo-bulk center cells are used as input to Palantir.

#### Pathway enrichment analysis

Gene set annotations used for pathway enrichment analyses are obtained using the msigdb_gsets function from the R package hypeR^64^. For human data, we used the "Homo sapiens","C5","BP" gene sets, while for mouse data, we used the "Mus musculus","C5","BP" gene sets. Pathway enrichment was calculated using Fisher’s exact test.

#### Tracking nucleosome binding dynamics across human hematopoiesis

To characterize nucleosome reorganization during human hematopoiesis, we implemented a custom script for automatic tracking of nucleosome footprints. Given any lineage of interest, we ordered all pseudo-bulks in the lineage by pseudo-time. Due to the sparsity of data, we applied a sliding pseudo-time window of 10 pseudo-bulks and combined the data in each window before footprinting. Nucleosome footprints (100 bp scale footprints) were then called at each pseudo-time point. Next, we aggregated the data from all pseudo-bulks and called nucleosome footprints. The aggregate nucleosome footprint centers were used as rough mapping of the position for each nucleosome. For fine-mapping of nucleosome position, we defined a window of 100 bp in diameter centered at the aggregate nucleosome footprint position. At each pseudo-time point, we found the position of maximum footprint score within this 100 bp window as the instantaneous position of the nucleosome. The instantaneous position as well as footprint intensity of the nucleosome were recorded for each nucleosome and pseudo-time point. Eventually, the position and footprint intensity of the same nucleosome were compared across pseudo-time to analyze sliding / binding / eviction of the nucleosome.

#### Characterizing age-related intra-cCRE dynamics

##### Data preprocessing

Cells with fraction of reads in peaks (FRIP) < 0.3 and depth < 300 were first removed. Additionally, we used ArchR^65^ to calculate doublet scores for each single cell and removed cells with top 5% doublet scores. The remaining cells were then processed with the Seurat package^66^. Cells were embedded into lower dimensional space using latent semantic indexing (LSI)^67^ and then clustered. Seurat clusters corresponding to HSCs were selected for pseudo-bulking and downstream differential testing. Cells with the “LinNeg” FACS sort label were excluded for HSC-specific analyses. To identify representative cell states, we used SEACells^38^ to identify 30 representative cell states across HSCs. The representative cells are used as centers to form pseudo-bulks. Each pseudo-bulk is generated by serially including nearest neighbor cells from the center cell in an order of increasing distance until we reach a total of 5 million reads.

##### Differential testing

Differential RNA testing was performed using DESeq2^68^. We first quantified total RNA read counts for each gene in each pseudo-bulk, and used DESeq2 to identify significant differential genes with age as the covariate. Additionally, we applied the TF habitation model to all cCREs and then segmented cCREs into sub-cCREs using the method mentioned above. After filtering out low signal sub-cCREs (< 0.3 activity), we performed differential cCRE and sub-cCRE testing with two-tailed unequal variance t-test (Welch’s test) using the cCRE accessibility and sub-cCRE activity, respectively.

##### Motif score analysis

Motif scores were calculated using chromVAR^49^. Unlike the standard practice using the cCRE-by-sample count matrix, here we used sub-cCRE-by-pseudo-bulk activity matrix. Only sub-cCREs with differential sub-cCRE FDR < 0.1 as well as differential cCRE FDR > 0.1 are included for motif scoring. We performed differential motif score analysis using two-tailed unequal variance t-test (Welch’s test) between young and old age groups.

## Data Availability

Additional data such as pre-trained machine learning models and pre-computed Tn5 bias tracks can be accessed on Zenodo at https://zenodo.org/record/7121027#.ZCLo0ezMI8M. Interactive visualization using Shinyapps can be found at https://buenrostrolab.shinyapps.io/ACAMShiny/ (human bone marrow) and https://buenrostrolab.shinyapps.io/aging/ (mouse HSC aging). Raw and processed sequencing data can be found on Gene Expression Omnibus (GEO) with the accession number GSE216464.

## Code Availability

All code used in this study, including tutorial for running PRINT, can be found at https://github.com/HYsxe/PRINT.

## Acknowledgements

We thank members of the Buenrostro lab and the Wagers Lab for useful discussions and critical assessment of this work. J.D.B. and the Buenrostro lab acknowledge support from the Gene Regulation Observatory at the Broad Institute of MIT & Harvard, the Chan Zuckerberg Initiative, the NHGRI IGVF consortium (UM1 HG011986) and the NIH New Innovator Award (DP2 HL151353). A.J.W. and the Wagers Lab acknowledge support by grants from NIH (DP1 OD025432) and the Glenn Foundation for Medical Research (to A.J.W.) and NIH F32 AG071208 to H.K. We thank J. LaVecchio N. Kheramand at the HSCI/HSCRB Flow Core for assistance with flow cytometry and FACS. We thank A. Brack and C. Epstein for assistance in data generation.

## Author contributions

Y. Hu led all computational developments and analyses described in this work with contributions from S. Ma, V. Kartha, R. Zhang, A. Meliki, A. Castillo, N. Durand, E. Mattei, and N. Shoresh. S. Ma, F. Duarte, M. Horlbeck, R. Shrestha, A. Labade, H. Kletzien, L. J. Anderson, T. Tay, and A. S. Earl generated the data with supervision by C. B. Epstein, A. Wagers, and J. Buenrostro. J. Buenrostro supervised all aspects of this work. Y. Hu and J. Buenrostro wrote the manuscript with input from all authors.

## References

1. Shlyueva, D., Stampfel, G. & Stark, A. Transcriptional enhancers: from properties to genome-wide predictions. Nat. Rev. Genet. 15, 272–286 (2014).

2. Gasperini, M., Tome, J. M. & Shendure, J. Towards a comprehensive catalogue of validated and target-linked human enhancers. Nat. Rev. Genet. 21, 292–310 (2020).

3. François Spitz & Eileen E. M. Furlong. Transcription factors: from enhancer binding to developmental control. Nat. Rev. Genet. (2012) doi:10.1038/nrg3207.

4. Segal, E. & Widom, J. From DNA sequence to transcriptional behaviour: a quantitative approach. Nat. Rev. Genet. 10, 443–456 (2009).

5. Calo, E. & Wysocka, J. Modification of enhancer chromatin: what, how, and why? Mol. Cell 49, 825–837 (2013).

6. ENCODE Project Consortium et al. Expanded encyclopaedias of DNA elements in the human and mouse genomes. Nature 583, 699–710 (2020).

7. de Dieuleveult, M. et al. Genome-wide nucleosome specificity and function of chromatin remodellers in ES cells. Nature 530, 113–116 (2016).

8. Lai, W. K. M. & Pugh, B. F. Understanding nucleosome dynamics and their links to gene expression and DNA replication. Nat. Rev. Mol. Cell Biol. 18, 548–562 (2017).

9. Jain, S. S. & Tullius, T. D. Footprinting protein-DNA complexes using the hydroxyl radical. Nat. Protoc. 3, 1092–1100 (2008).

10. Galas, D. J. & Schmitz, A. DNAse footprinting: a simple method for the detection of protein-DNA binding specificity. Nucleic Acids Res. 5, 3157–3170 (1978).

11. Hesselberth, J. R. et al. Global mapping of protein-DNA interactions in vivo by digital genomic footprinting. Nat. Methods 6, 283–289 (2009).

12. Neph, S. et al. An expansive human regulatory lexicon encoded in transcription factor footprints. Nature 489, 83–90 (2012).

13. Buenrostro, J. D., Giresi, P. G., Zaba, L. C., Chang, H. Y. & Greenleaf, W. J. Transposition of native chromatin for fast and sensitive epigenomic profiling of open chromatin, DNA-binding proteins and nucleosome position. Nat. Methods 10, 1213–1218 (2013).

14. Zentner, G. E. & Henikoff, S. High-resolution digital profiling of the epigenome. Nat. Rev. Genet. 15, 814–827 (2014).

15. Degner, J. F. et al. DNase I sensitivity QTLs are a major determinant of human expression variation. Nature 482, 390–394 (2012).

16. Stergachis, A. B. et al. Developmental fate and cellular maturity encoded in human regulatory DNA landscapes. Cell 154, 888–903 (2013).

17. Neph, S. et al. Circuitry and dynamics of human transcription factor regulatory networks. Cell 150, 1274– 1286 (2012).

18. Stergachis, A. B. et al. Conservation of trans-acting circuitry during mammalian regulatory evolution. Nature 515, 365–370 (2014).

19. He, H. H. et al. Refined DNase-seq protocol and data analysis reveals intrinsic bias in transcription factor footprint identification. Nat. Methods 11, 73–78 (2014).

20. Adey, A. et al. Rapid, low-input, low-bias construction of shotgun fragment libraries by high-density in vitro transposition. Genome Biol. 11, R119 (2010).

21. Bentsen, M. et al. ATAC-seq footprinting unravels kinetics of transcription factor binding during zygotic genome activation. Nat. Commun. 11, 4267 (2020).

22. Li, Z. et al. Identification of transcription factor binding sites using ATAC-seq. Genome Biol. 20, 45 (2019).

23. Schep, A. N. et al. Structured nucleosome fingerprints enable high-resolution mapping of chromatin architecture within regulatory regions. Genome Res. 25, 1757–1770 (2015).

24. Voong, L. N. et al. Insights into Nucleosome Organization in Mouse Embryonic Stem Cells through Chemical Mapping. Cell 167, 1555–1570.e15 (2016).

25. Puig, R. R., Boddie, P., Khan, A., Castro-Mondragon, J. A. & Mathelier, A. UniBind: maps of high-confidence direct TF-DNA interactions across nine species. BMC Genomics 22, 482 (2021).

26. Ma, S. et al. Chromatin Potential Identified by Shared Single-Cell Profiling of RNA and Chromatin. Cell 183, 1103–1116.e20 (2020).

27. Niccoli, T. & Partridge, L. Ageing as a risk factor for disease. Curr. Biol. 22, R741–52 (2012).

28. Pal, S. & Tyler, J. K. Epigenetics and aging. Sci Adv 2, e1600584 (2016).

29. Zhang, W., Qu, J., Liu, G.-H. & Belmonte, J. C. I. The ageing epigenome and its rejuvenation. Nat. Rev. Mol. Cell Biol. 21, 137–150 (2020).

30. Riedel, C. G. et al. DAF-16 employs the chromatin remodeller SWI/SNF to promote stress resistance and longevity. Nat. Cell Biol. 15, 491–501 (2013).

31. Morrison, S. J., Wandycz, A. M., Akashi, K., Globerson, A. & Weissman, I. L. The aging of hematopoietic stem cells. Nat. Med. 2, 1011–1016 (1996).

32. Chambers, S. M. et al. Aging hematopoietic stem cells decline in function and exhibit epigenetic dysregulation. PLoS Biol. 5, e201 (2007).

33. Rossi, D. J., et al. Cell intrinsic alterations underlie hematopoietic stem cell aging. Proc. Natl. Acad. Sci. U. S. 102, 9194–9199 (2005).

33. Evans, M. A. & Walsh, K. Clonal Hematopoiesis, Somatic Mosaicism, and Age-Associated Disease. Physiol. Rev. (2022) doi:10.1152/physrev.00004.2022.

34. Jaiswal, S. & Ebert, B. L. Clonal hematopoiesis in human aging and disease. Science 366, (2019).

35. Sun, D., et al. Epigenomic Profiling of Young and Aged HSCs Reveals Concerted Changes during Aging that Reinforce Self-Renewal. Cell Stem Cell vol. 14 673–688 Preprint at https://doi.org/10.1016/j.stem.2014.03.002 (2014).

36. Mansell, E. et al. Mitochondrial Potentiation Ameliorates Age-Related Heterogeneity in Hematopoietic Stem Cell Function. Cell Stem Cell 28, 241–256.e6 (2021).

37. Persad, S. et al. SEACells: Inference of transcriptional and epigenomic cellular states from single-cell genomics data. bioRxiv 2022.04.02.486748 (2022) doi:10.1101/2022.04.02.486748.

38. Hsu, A.-L., Murphy, C. T. & Kenyon, C. Regulation of aging and age-related disease by DAF-16 and heat-shock factor. Science 300, 1142–1145 (2003).

39. Labbadia, J. et al. Mitochondrial Stress Restores the Heat Shock Response and Prevents Proteostasis Collapse during Aging. Cell Rep. 21, 1481–1494 (2017).

40. Moll, L. et al. The insulin/IGF signaling cascade modulates SUMOylation to regulate aging and proteostasis in Caenorhabditis elegans. Elife 7, (2018).

41. Ho, T. T. et al. Autophagy maintains the metabolism and function of young and old stem cells. Nature 543, 205–210 (2017).

42. Itokawa, N. et al. Epigenetic traits inscribed in chromatin accessibility in aged hematopoietic stem cells. Nat. Commun. 13, 2691 (2022).

43. Vierstra, J. et al. Global reference mapping of human transcription factor footprints. Nature 583, 729–736 (2020).

44. Bergsland, M. et al. Sequentially acting Sox transcription factors in neural lineage development. Genes Dev. 25, 2453–2464 (2011).

45. Jing, H. et al. Exchange of GATA factors mediates transitions in looped chromatin organization at a developmentally regulated gene locus. Mol. Cell 29, 232–242 (2008).

46. Zhang, Y. et al. Model-based analysis of ChIP-seq (MACS). Genome Biol. 9, R137 (2008).

47. Heinz, S., et al. Simple Combinations of Lineage-Determining Transcription Factors Prime cis-Regulatory Elements Required for Macrophage and B Cell Identities. Molecular Cell vol. 38 576–589 Preprint at https://doi.org/10.1016/j.molcel.2010.05.004 (2010).

48. Schep, A. N., Wu, B., Buenrostro, J. D. & Greenleaf, W. J. chromVAR: inferring transcription-factor-associated accessibility from single-cell epigenomic data. Nat. Methods 14, 975–978 (2017).

49. Bravo González-Blas, C., et al. cisTopic: cis-regulatory topic modeling on single-cell ATAC-seq data. Nat. Methods 16, 397–400 (2019).

50. McInnes, L., Healy, J., Saul, N. & Großberger, L. UMAP: Uniform Manifold Approximation and Projection. Journal of Open Source Software vol. 3 861 Preprint at https://doi.org/10.21105/joss.00861 (2018).

51. Stuart, T. et al. Comprehensive Integration of Single-Cell Data. Cell 177, 1888–1902.e21 (2019).

52. Nair & Hinton. Rectified linear units improve restricted boltzmann machines. Icml (2010).

53. Chollet, F. & Others. Keras. https://keras.io (2015).

54. Kingma, D. P. & Ba, J. Adam: A Method for Stochastic Optimization. Preprint at https://doi.org/10.48550/ARXIV.1412.6980 (2014).

55. Kent, W. J. The Human Genome Browser at UCSC. Genome Research vol. 12 996–1006 Preprint at https://doi.org/10.1101/gr.229102. (2002).

56. Schep, A. motifmatchr: Fast Motif Matching in R. (2022).

57. Li, Z. et al. Chromatin-accessibility estimation from single-cell ATAC data with scOpen. bioRxiv 865931 (2021) doi:10.1101/865931.

58. Mei, S. et al. Cistrome Data Browser: a data portal for ChIP-seq and chromatin accessibility data in human and mouse. Nucleic Acids Res. 45, D658–D662 (2017).

59. Hahne, F. & Ivanek, R. Visualizing Genomic Data Using Gviz and Bioconductor. Methods Mol. Biol. 1418, 335–351 (2016).

60. Martín Abadi, et al. TensorFlow: Large-Scale Machine Learning on Heterogeneous Systems. Preprint at https://www.tensorflow.org/ (2015).

61. Crane, E. et al. Condensin-driven remodelling of X chromosome topology during dosage compensation. Nature 523, 240–244 (2015).

62. Setty, M. et al. Characterization of cell fate probabilities in single-cell data with Palantir. Nat. Biotechnol. 37, 451–460 (2019).

63. Federico, A. & Monti, S. hypeR: an R package for geneset enrichment workflows. Bioinformatics 36, 1307– 1308 (2020).

64. Granja, J. M. et al. ArchR is a scalable software package for integrative single-cell chromatin accessibility analysis. Nat. Genet. 53, 403–411 (2021).

65. Satija, R., Farrell, J. A., Gennert, D., Schier, A. F. & Regev, A. Spatial reconstruction of single-cell gene expression data. Nat. Biotechnol. 33, 495–502 (2015).

66. Deerwester, Dumais, Furnas, Landauer & Harshman. Indexing by latent semantic analysis. J. Am. Soc. Inf. Sci.

67. Love, M. I., Huber, W. & Anders, S. Moderated estimation of fold change and dispersion for RNA-seq data with DESeq2. Genome Biol. 15, 550 (2014).

