## Supplementary Information for "Single-cell multi-scale footprinting reveals the modular organization of DNA regulatory elements"

### Calculation of footprint scores

Framework for statistical testing

For any specific position in the cCRE, we define a center footprint region and a flanking region (Fig. 1**e**). We then calculate a footprinting score using statistical testing. The test statistic $\lambda$ is the ratio of total Tn5 insertions in the footprint region divided by the total Tn5 insertions in the footprint and flanking regions combined.

$$\lambda = \frac{\sum_{i \in A_{footprint}} x_i}{\sum_{i \in A_{flank}} x_i + \sum_{i \in A_{footprint}} x_i}$$

Here, $x_i$ is the number of Tn5 insertions at position $i$. $A_{flank}$ and $A_{footprint}$ are the sets of position indices in the flanking and footprint regions, respectively. The goal is to estimate the background distribution of $\lambda$ when no protein is bound, and then compare the observed value of $\lambda$ to its background distribution. If the position of interest is bound by protein, the observed $\lambda$ should be significantly lower than the background distribution. Hence, we can calculate a *p*-value to represent the significance of such deviation.

It is worth mentioning that here we decide to perform two tests on each side (i.e., center-vs-left and center-vs-right) and then keep the less significant *p*-value as the result. More specifically, we calculate the below $\lambda_{left}$ and $\lambda_{right}$.

$$\lambda_{left} = \frac{\sum_{i \in A_{footprint}} x_i}{\sum_{i \in A_{flankL}} x_i + \sum_{i \in A_{footprint}} x_i}, \lambda_{right} = \frac{\sum_{i \in A_{footprint}} x_i}{\sum_{i \in A_{flankR}} x_i + \sum_{i \in A_{footprint}} x_i}$$

$A_{flankL}$ and $A_{flankR}$ are the sets of position indices in the left and right flanking regions, respectively. The reason behind testing each side is to reduce false positive results. Consider the case illustrated below where one accessible cCRE is flanked by two nucleosomes:

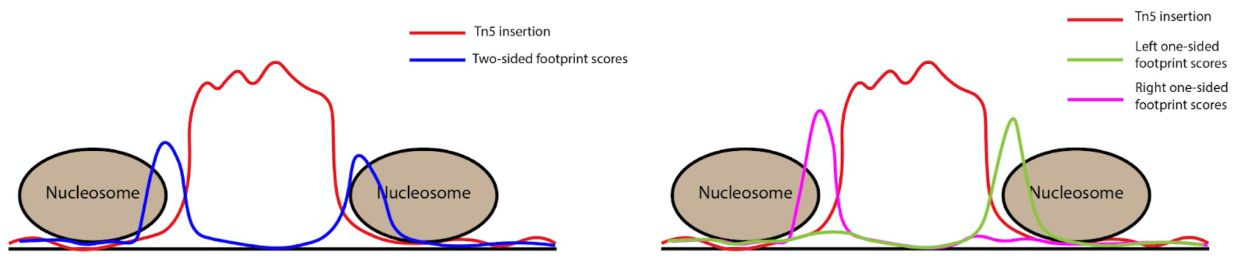


For positions at the edge of the accessible region, Tn5 insertion will be low in the center footprint region as well as one of the two flanking regions. High Tn5 insertion will be observed in the other flanking region. If we compare the footprint region with both flanks combined, we might detect false positive signal since $\lambda$ can still be lower than expected. On the contrary, performing two tests on each side and keeping the less significant result will efficiently solve such issues.

Modeling the background distribution

The BAC naked DNA Tn5 insertion data was used to estimate the background distribution of $\lambda$. We reason that for each flanking side, the background distribution of $\lambda$ should be determined by center and flanking Tn5 bias, as well as the total number of reads in center and flank regions. Suppose $b$ is the vector of predicted Tn5 bias (i.e., predicted bias at position $i$ is $b_i$), we have

$$b_{left} = \sum_{\in A_{flankL}} b_i,\ b_{center} = \sum_{\in A_{center}} b_i,\ b_{right} = \sum_{\in A_{flankR}} b_i$$

$$c_{left} = \sum_{i\in A_{flankL} \cup A_{center}} x_i\,, c_{right} = \sum_{i\in A_{flankR} \cup A_{center}} x_i$$

$b_{left}$, $b_{center}$, and $b_{right}$ are total biases in the left flanking, center footprint, and right flanking regions, respectively. $c_{left}$, and $c_{right}$ are coverage for the left and right side testing, respectively. We then aim to model the below distributions

$$\lambda_{left} \sim F_{left}(b_{left}, b_{center}, c_{left})$$

$$\lambda_{right} \sim F_{right}(b_{right}, b_{center}, c_{right})$$

To this end, the most straightforward approach would be to use the BAC naked DNA data as a lookup table. Suppose we want to compute the footprint score for a specific position (here referred to as “foreground”) in an ATAC-Seq dataset. We compute the center bias, flanking bias, as well as coverage for the foreground observation. Next, we search the BAC naked DNA data to find the 500 nearest neighbor observations in the (center bias, flanking bias, coverage) 3-dimensional space. We compute the $\lambda$ for these background observations and denote as $\lambda_{bg}$. The distribution of $\lambda_{bg}$ is then used as the background distribution for the foreground observed ratio $\lambda_{obs}$ and a *p*-value is computed using *z*-test. To make sure the KNN matching weighs the three features equally, coverage values are first log10 transformed and then all three features are standardized before KNN matching. Additionally, to make sure we cover a wide range of coverage values, the BAC dataset was down-sampled to 100%, 50%, 20%, 10%, 5%, 2%, and 1% and then pooled before KNN matching.

In practice, performing KNN matching for each foreground observation is extremely time-consuming. Therefore, we instead train a neural network dispersion model to learn the below relationship:

$$(\mu_{left}, SD_{left}, \mu_{right}, SD_{right}) \sim F(b_{left}, b_{right}, b_{center}, c_{left}, c_{right})$$

We randomly sample 100,000 observations from the BAC dataset. For each of them, we match 500 nearest neighbor observations in the BAC dataset and compute the distribution of $\lambda_{bg}$. Then these 100,000 observations along with their background distribution are used to train the dispersion model.

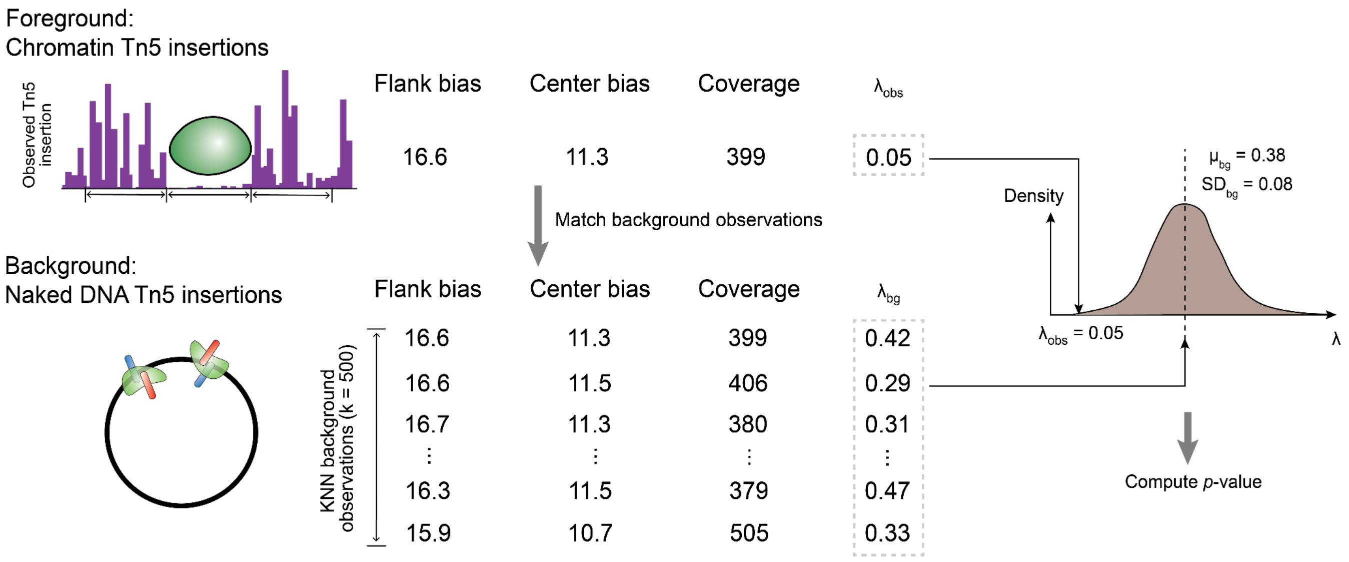


### Multi-scale footprinting

There are two main motivations behind using multi-scale footprinting for object detection. The first one is that some DNA-binding factors do not leave footprints on their own, potentially due to weak or transient binding. Therefore, we can only infer the binding of such factors through the binding and positioning of nearby objects such as nucleosomes. In these cases, using only footprints detected at the scale corresponding to the size of the factor itself will lead to false negative results (e.g., only using the 40 bp scale footprint to detect YY1 binding).

The second motivation is signal bleed through across scales. We realized that the footprint signal of an object can bleed through into lower scales, potentially due to non-linear impact on Tn5 insertion by object binding. For instance, examine the example below：

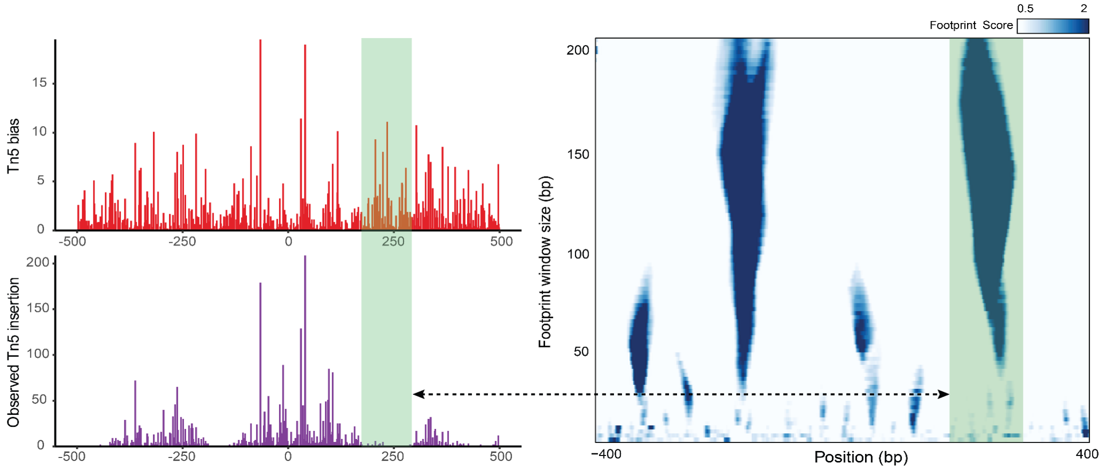

$$M_{i,j} = cor(S_i, S_j)$$

The highlighted boxes in the left and right plots represent the same genomic region. We can see that this region shows footprint signals from around 40 up to 200 bp scales, representing a bound nucleosome. Although the nucleosome-bound region is around 140 bp in diameter, the footprint signal is not limited to 140 bp scale, but bleeds through to a much lower spatial scale. In the case shown above, this is because nucleosome binding does not affect Tn5 insertion at all positions equally. Positions with very high Tn5 bias, shown as "spikes"' in the Tn5 bias track, are affected most dramatically. In the highlighted region, several positions have Tn5 bias spikes, and are thus expected by the model to have a high center / (center + flank) ratio of Tn5 insertion. However, in the observed insertion track, due to the binding of the nucleosome, Tn5 insertion at these locations are suppressed to a low baseline level, similar to flaking regions. Therefore, the observed center / (center + flank) ratio is lower than the expected, and the model will return significant footprint signals at such positions. In essence, the above results from the unequal impact of the bound object on Tn5 insertion. In other words, if insertion at all base pairs in the bound regions are reduced by the same scaling factor, including positions with Tn5 bias spikes, then most of the bleed-through will no longer appear.

As a result, if we detect TF binding only by examining footprint signals at the scale matching the size of TFs, such bleed-through will result in false positive signals. The bleed-through effect is theoretically not specific to our footprinting method. For any method that defines a footprint window and calculates deviation of observed Tn5 insertion in the footprint window from the expected level, this could be a potential source of false positive signal. Positions with Tn5 bias spikes but bound by nucleosomes will tend to have high expected but low observed cutting, and could show up as a false positive footprint at the scale of TFs.

Given the above, we realize that trying to detect objects using footprint scores at a single scale and a single location is under-powered. Leveraging footprint signals across scales and positions eventually became our option for accurate object detection and tracking.

#### Segmentation of sub-cCREs

For segmentation of cCREs into sub-cCRE, we first divide each cCRE into 10 bp intervals as candidate TF binding sites, and then calculate a boundary score for each site. The boundary score evaluates whether a certain site should be assigned as a sub-cCRE boundary. Suppose for a specific cCRE we have $k$ candidate TF binding sites and data from $n$ pseudo-bulks. We start with a $k$-by-$n$ matrix $S$ of TF habitation scores. We use $S$ to calculate a $k$-by-$k$ pairwise correlation matrix $M$

$$M_{i,j} = cor(S_i, S_j)$$

Then for each TF site, we define a local region with radius $r$, and calculate 3 scores representing TF co-binding upstream, across, and downstream of the current site of interest, respectively. Therefore, we have

$$ScoreA = \sum_{i=max(1,x-r+1)}^{x} \sum_{j=max(1,x-r+1)}^{x} \frac{M_{i,j}}{min(x,r)^2}$$

$$ScoreB = \sum_{i=max(1,x-r+1)}^{x} \sum_{j=x+1}^{min(k,x+r)} \frac{M_{i,j}}{min(x,r) \times min(r,k-x)}$$

$$ScoreC = \sum_{i=x+1}^{\min(k,x+r)} \sum_{j=x+1}^{min(k,x+r)} \frac{M_{i,j}}{min(r,k-x)^2}$$

Combining these 3 scores, we get the boundary score:

$$boundary\ score = max(ScoreA, ScoreC) - ScoreB$$

See the below schematic

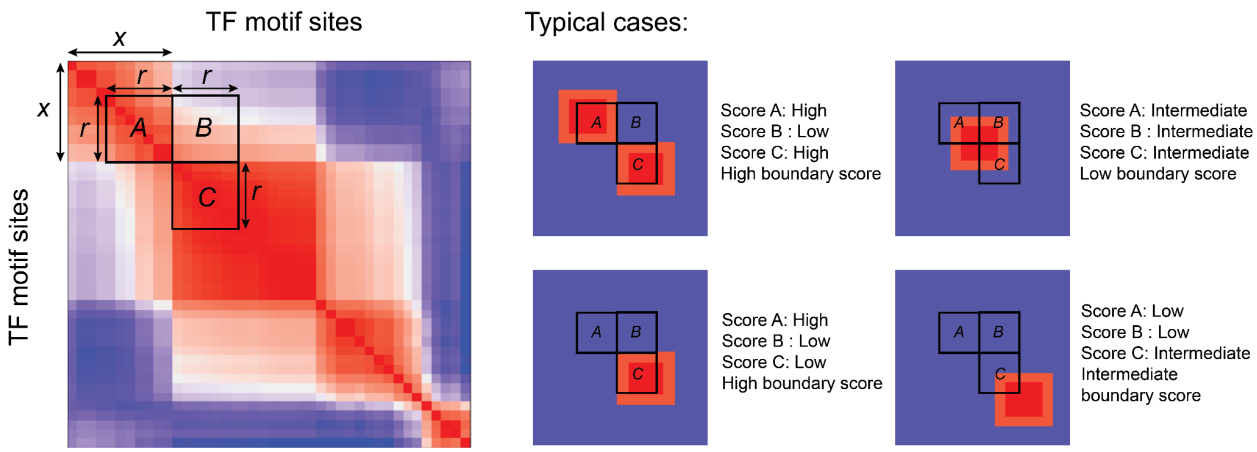


This is similar to calculating insulation scores for segmenting topologically associating domains (TADs)[1]. Our boundary score is essentially applying a pattern matching kernel to each position along the diagonal of the correlation map. It serves to detect positions with high correlation on either side of the TF binding site but not across it. As the pattern matching kernel slides along the diagonal, we will see gradual rise and fall of the boundary score. We decided to first smooth the boundary scores and assign local maximum boundary score positions as sub-cCRE boundaries. Eventually, the boundary positions are mapped back to the original genomic positions in the cCRE, resulting in segmentation as we see in the top color bar of Fig. 3**b**.


1. Crane, E. *et al.* Condensin-driven remodelling of X chromosome topology during dosage compensation. *Nature* **523**, 240–244 (2015).
