## Supplementary figures and images for "Single-cell multi-scale footprinting reveals the modular organization of DNA regulatory elements"

### Extended Data File 1

##### FACS plots (young male mice, n=10):

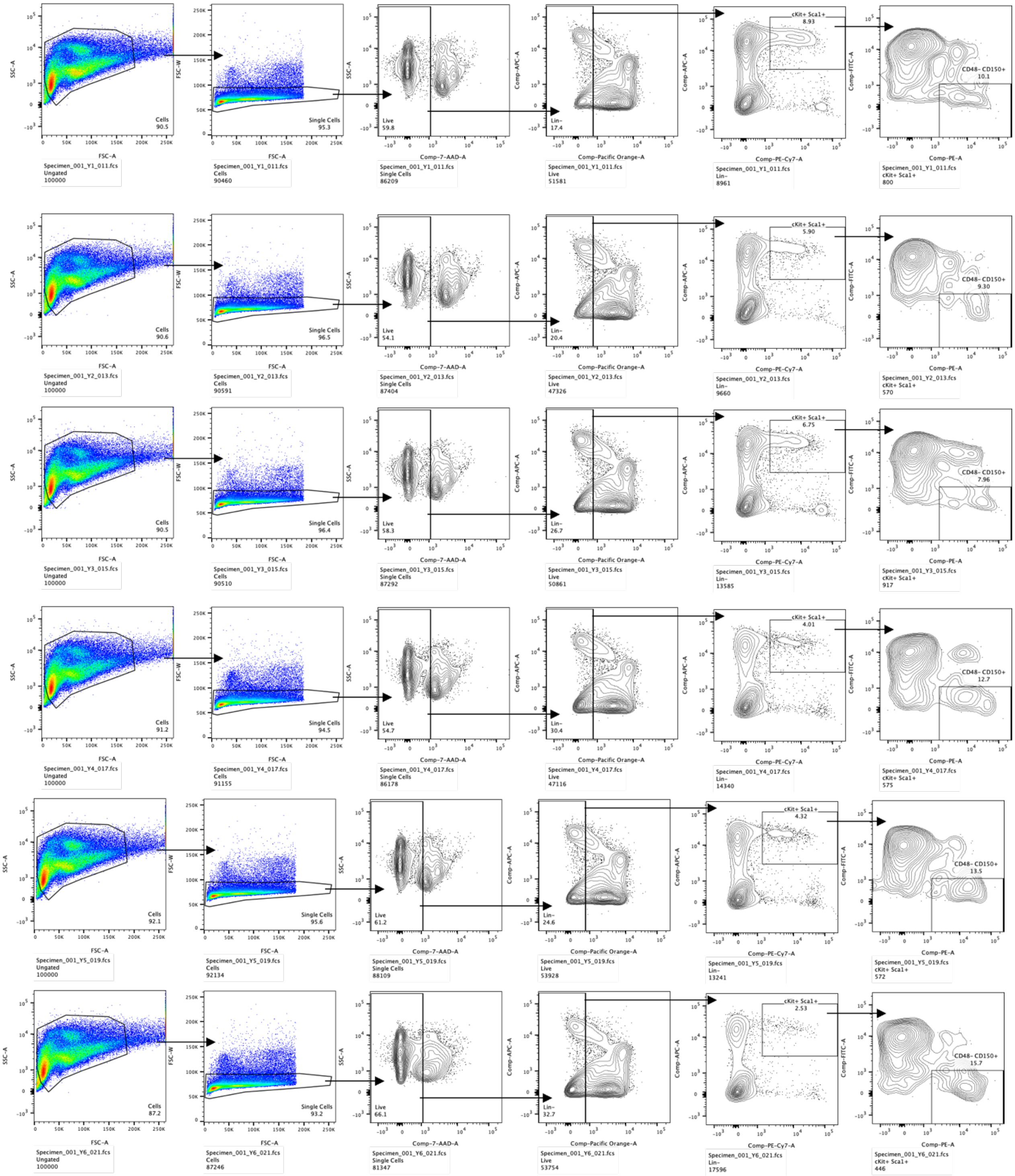

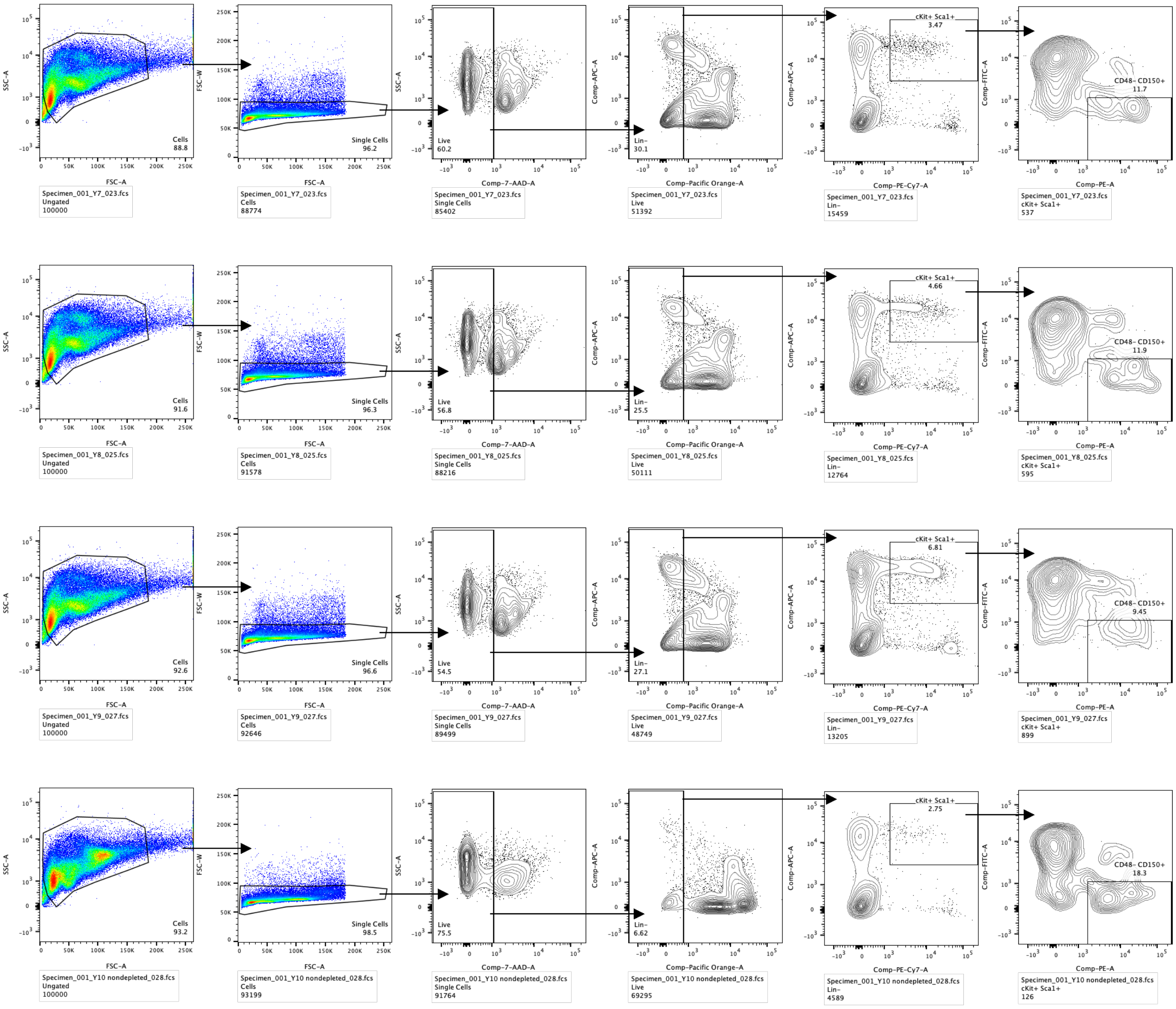


#### FACS plots (aged male mice, n=5):

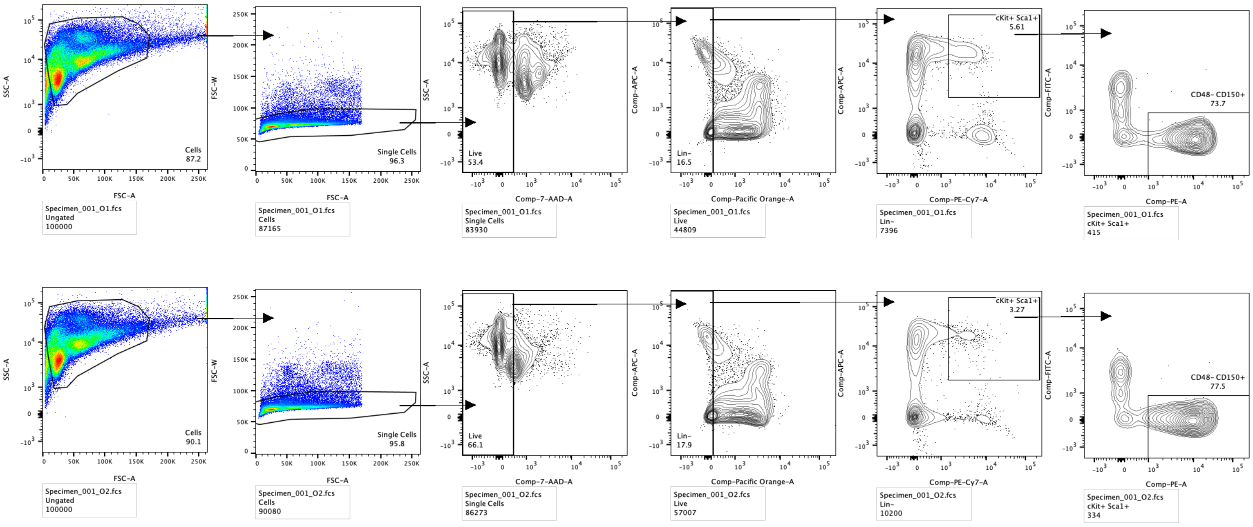

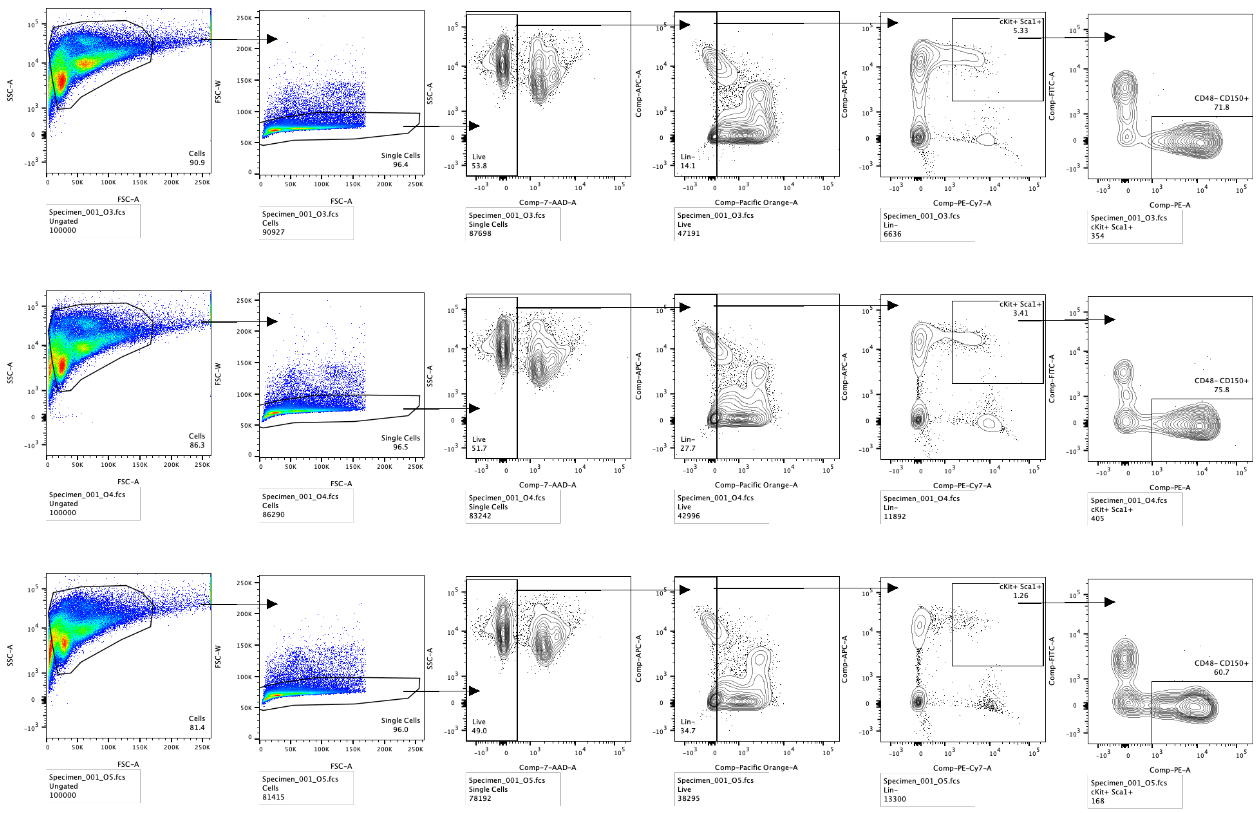


#### Purity/Resort:

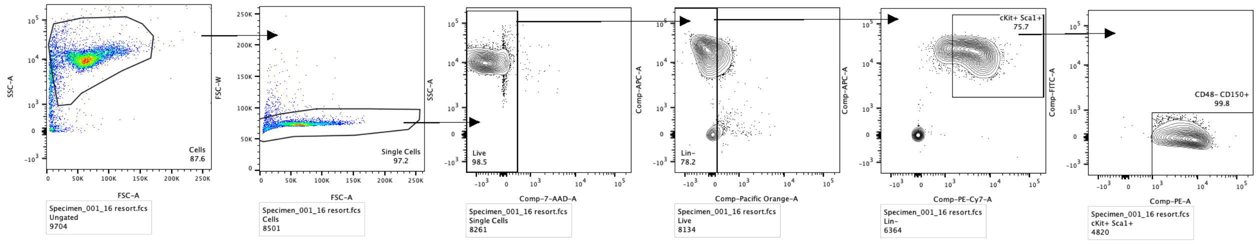
